# Atf3 links loss of epithelial polarity to defects in cell differentiation and cytoarchitecture

**DOI:** 10.1101/212969

**Authors:** Colin D. Donohoe, Gábor Csordás, Andreia Correia, Marek Jindra, Corinna Klein, Bianca Habermann, Mirka Uhlirova

## Abstract

Interplay between apicobasal cell polarity modules and the cytoskeleton is critical for differentiation and integrity of epithelia. However, this coordination is poorly understood at the level of gene regulation by transcription factors. Here, we establish the *Drosophila activating transcription factor 3* (*atf3*) as a cell polarity response gene acting downstream of the membrane-associated Scribble polarity complex. Loss of the tumor suppressors Scribble or Dlg1 induces *atf3* expression via aPKC but independent of Jun-N-terminal kinase (JNK) signaling. Strikingly, removal of Atf3 from Dlg1 deficient cells restores polarized cytoarchitecture, levels and distribution of endosomal trafficking machinery, and differentiation. Conversely, excess Atf3 alters microtubule network, vesicular trafficking and the partition of polarity proteins along the apicobasal axis. Genomic and genetic approaches implicate Atf3 as a regulator of cytoskeleton organization and function, and identify *Lamin C* as one of its *bona fide* target genes. By affecting structural features and cell morphology, Atf3 functions in a manner distinct from other transcription factors operating downstream of disrupted cell polarity.

**Author summary:** Epithelial cells form sheets and line both the outside and inside of our body. Their proper development and function require the asymmetric distribution of cellular components from the top to the bottom, known as apicobasal polarization. As loss of polarity hallmarks a majority of cancers in humans understanding how epithelia respond to a collapse of the apicobasal axis is of great interest. Here, we show that in the fruit fly *Drosophila melanogaster*, the breakdown of epithelial polarity engages Activating transcription factor 3 (Atf3), a protein that directly binds the DNA and regulates gene expression. We demonstrate that many of the pathological consequences of disturbed polarity require Atf3, as its loss in this context results in normalization of cellular architecture, vesicle trafficking and differentiation. Using unbiased genome-wide approaches we identify the genetic program controlled by Atf3 and experimentally verify select candidates. Given the evolutionary conservation of Atf3 between flies and man, we believe that our findings in the *Drosophila* model will contribute to a better understanding of diseases stemming from compromised epithelial polarity.

## Introduction

Epithelia are sheets of highly polarized cells that represent the defining tissue type of metazoans. With their capacity to achieve various shapes and serve as selective barriers, epithelial tissues play vital roles in morphogenesis, tissue differentiation and compartmentalization, and intercellular signaling. The integrity and function of epithelia rely on the interplay between key polarity determinants and a highly ordered yet dynamic cytoskeleton, which ensures tissue plasticity and the asymmetric distribution of cellular components along the apicobasal polarity axis [1].

Genetic studies have established a network of evolutionarily conserved signaling pathways and effector molecules that govern the organization of epithelial cellular architecture. In particular, the basic leucine zipper (bZIP) transcription factors, including Jun, Fos, and Atf3, are important regulators of epithelial function from the fruit fly *Drosophila* to mammals [2-6]. During *Drosophila* development, expression of *atf3* is dynamic and under tight temporal constraints. Ectopic Atf3 activity in larval epidermal cells (LECs) disturbs epithelial morphogenesis of the adult abdomen, stemming from perturbed cytoskeleton dynamics and increased cell adhesion which prevent normal LEC extrusion [6].

The importance of finely tuned Atf3 expression during *Drosophila* development corresponds with the role of mammalian ATF3 as a stress-response gene regulated at the level of mRNA expression by various stimuli, including genotoxic radiation, wounding, cytokines, nutrient deprivation, Toll signaling, oncogenes and inhibition of calcineurin-NFAT signaling [7-9]. Independent transcriptome analyses of *Drosophila* epithelia have shown deregulated *atf3* expression in wing imaginal discs lacking the conserved neoplastic tumor suppressor genes *scribble* (*scrib*) or *discs large 1* (*dlg1*) encoding components of the Scribble polarity module [10], and in *ras*^*V12*^*scrib*^−^ tumors in the eye/antennal imaginal disc (EAD) [11-13]. These results point to loss of epithelial integrity as a novel trigger of *atf3* expression and are congruent with studies linking Atf3 to processes involving transient controlled epithelial depolarization during morphogenesis and wound healing [14-16] as well as to pathological disturbances in polarity that characterize chronic wounds and tumorigenesis [5,9,17-21]. However, which polarity cues induce *atf3* expression and how Atf3 activity contributes to phenotypes associated with loss of polarity have yet to be determined.

In this study, we establish that loss of the Scrib polarity module is sufficient to increase the levels and activity of Atf3 via aPKC signaling. Increased Atf3 activity drives major phenotypic attributes of the Dlg1 deficiency as abnormal distribution of polarity proteins and differentiation defects in *dlg1* mutant epithelial clones can be alleviated by removal of Atf3. Chromatin immunoprecipitation followed by high-throughput sequencing further revealed that Atf3 target genes are enriched for roles in cytoskeletal organization and dynamics. Thus, Atf3 links defects in the Scrib polarity module with gene dysregulation and subsequent perturbations in cellular morphology and differentiation.

## Results

### *atf3* is a cell-polarity response gene activated by aPKC but not JNK signaling

Previous transcriptome profiling by our group and others [11-13] has suggested that disturbed cell polarity leads to upregulation of *atf3* expression. Consistently, qRT-PCR from *scrib*^*1*^ homozygous mutant larvae and adult heads bearing *dlg1*^*G0342*^ homozygous mutant clones showed increased levels of *atf3* mRNA (Fig 1A), thus confirming induction of *atf3* transcription upon depletion of the Scrib polarity module. To extend this evidence, we tested whether Scrib or Dlg1 deficiency impacts the levels of an Atf3::GFP fusion protein expressed from a recombineered BAC (*atf3*^*gBAC*^) that is sufficient to rescue the lethality of the *atf3*^*76*^ null mutants [22]. In control eye/antennal imaginal disc (EAD), Atf3 was enriched in differentiated photoreceptors of the eye primordium and a subset of peripodial cells of the antenna (Fig 1B). In wing imaginal discs, Atf3::GFP protein labeled the squamous cells of the peripodial epithelium and columnar cells of the outer ring that encircles the wing pouch (Fig 1C). Knockdown of *dlg1* (*en>dlg1*^*RNAi*^) or *scrib* (*en>scrib*^*RNAi*^) in the posterior compartment of the wing disc resulted in a marked increase of the Atf3::GFP signal in the columnar epithelium of the wing pouch (Fig 1D, 1E). Importantly, Atf3 was also upregulated in *dlg1*^*G0342*^ loss-of-function mutant clones induced in the larval EAD (Fig 1F, 1G and S1A-C Fig).

**Fig 1.**
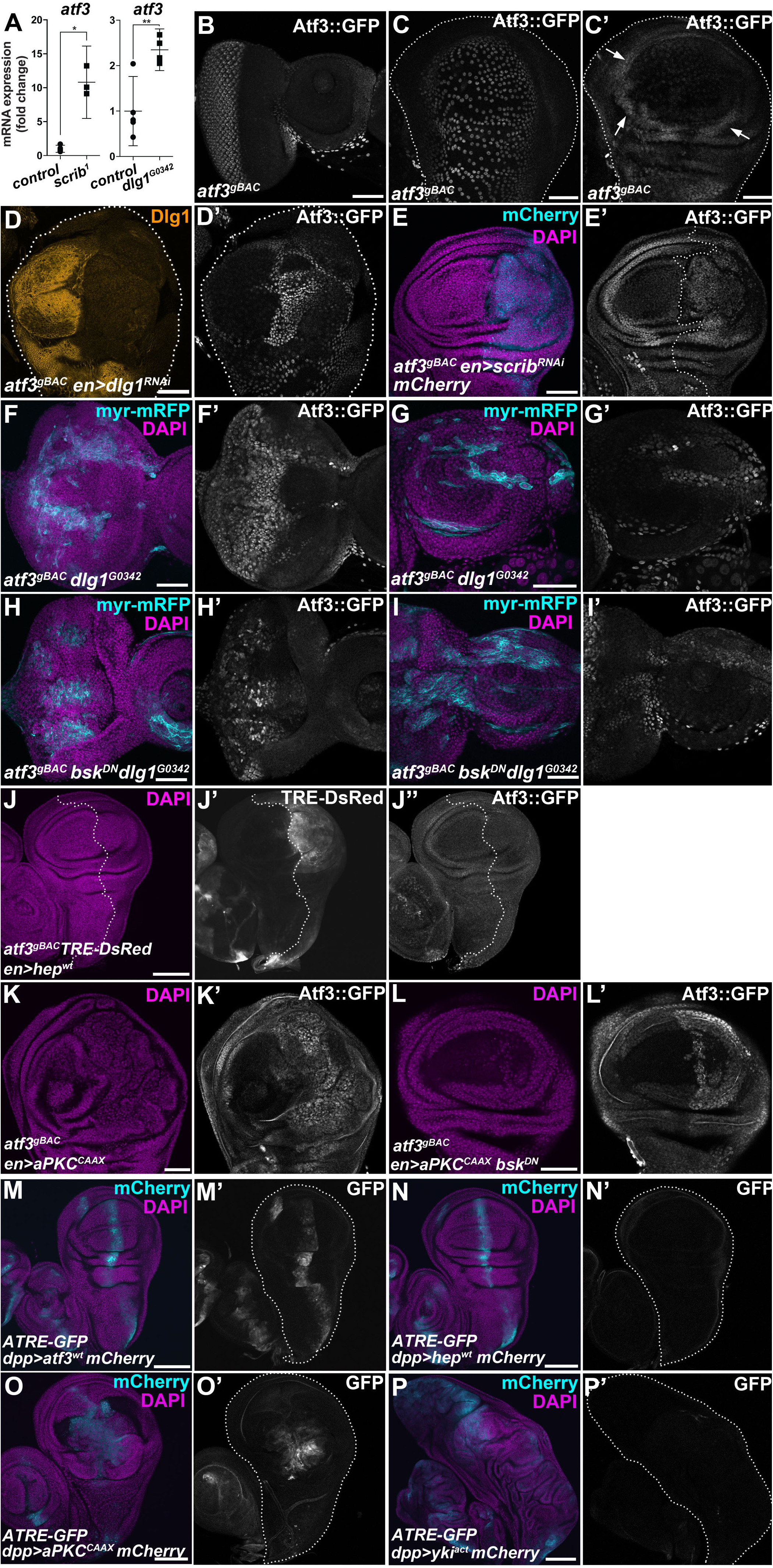
*atf3* is a polarity response gene. **(A)** Transcript levels of *atf3* increased in *scrib*^*1*^ mutant larvae and in adult heads bearing *dlg1*^*G0342*^ clones relative to control. qRT-PCR data are means of 5 and 3 biological replicates (*FRT82B* and *FRT82B scrib*^*1*^, respectively) and 5 and 4 biological replicates (*FRT19* and *FRT19 dlg1*^*G0342*^, respectively). Error bars depict 95% confidence interval; Unpaired Student’s t-tests assuming unequal variance were used to calculate p-values: *p=0.014, **p=0.0051. **(B-C)** Atf3::GFP fusion protein expressed from a recombineered BAC (*atf3*^*gBAC*^) labeled differentiated photoreceptors and a subset of peripodial cells in the control eye/antennal imaginal disc (B). In wing imaginal disc, peripodial cells (C) and columnar cells forming a ring surrounding the wing pouch (C’, arrows) showed Atf3::GFP enrichment. **(D-E)** Increase of Atf3::GFP signal appeared in the posterior compartment of the wing imaginal disc where Dlg1 (D-D’) or Scrib (E-E’) was depleted. **(F-I)** Clones in the EAD (myr-mRFP) deficient for Dlg1 expressed ectopic Atf3 within the eye (F’) and antenna (G’) part, which persisted when JNK signaling was blocked (*dlg1*^*G0342*^ *bsk*^*DN*^) (H’, I’). **(J)** Mild activation of JNK in the posterior compartment (J) of the wing imaginal disc (*en>hep*^*wt*^) induced TRE-DsRed (J’) but did not trigger expression of GFP-tagged Atf3 (J’’). **(K,L)** Atf3 protein levels increased following expression of a membrane tethered aPKC in the posterior compartment of the wing imaginal disc (*en>aPKC*^*CAAX*^) (K’). This induction was independent of JNK signaling (*en>aPKC*^*CAAX*^ *bsk*^*DN*^) (L’). **(M-P)** Driving Atf3 (M) or aPKC^CAAX^ (O) with *dpp-GAL4* (mCherry) induces an Atf3-responsive ATRE-GFP reporter (M’,O’), which is insensitive to Hep (N’) or activated Yki (P’). The wing discs (C-D, M-P) or the posterior wing disc compartments (E,J) are outlined with dotted lines. Micrographs B and F-I are projections of multiple confocal sections, while micrographs C-E and J-P are single slices. To allow for comparisons, micrographs M-P were acquired with the same microscope settings. Scale bars: 50 μm (B-I,K-L) and 100 μm (J,M-P).

Previous studies have shown that disruption of the Scrib module leads to activation of Jun-N-terminal kinase (JNK) signaling and to deregulation of the atypical protein kinase C (aPKC) [23-26]. To determine if either one of these two pathways is sufficient to induce Atf3 expression, we assessed both Atf3::GFP protein levels and the activity of a synthetic Atf3-responsive element (ATRE) reporter. The ATRE construct expresses GFP or RFP under the control of four concatenated genomic segments, each of which is 22 bp long and contains an Atf3 binding site identified through ChIP-seq in this study (S1 Table, Materials and Methods). Both *in vivo* and in cultured *Drosophila* S2 cells, Atf3 activated the ATRE reporter (Fig 1M’ and S2B, S2E Fig) but not its mutated version (mATRE) bearing three base pair substitutions in the Atf3 recognition sequence (S2D Fig and S1 Table). While activation of JNK signaling by expressing a wild type form of *Drosophila* JNKK (Hemipterous; Hep) induced a JNK-responsive TRE-DsRed reporter [27] (Fig 1J and S2G Fig), it failed to upregulate Atf3 expression when targeted to the posterior compartment of the wing imaginal disc (*en>hep*^*wt*^) (Fig 1J’’). Similarly, the ATRE-GFP reporter was insensitive to or only minimally activated by JNK signaling in the wing imaginal disc (*dpp>hep*^*wt*^) and S2 cells, respectively (Fig 1N and S2E Fig). Moreover, JNK signaling appeared dispensable for Atf3 induction caused by loss of polarity, as elevated Atf3::GFP signal persisted in *dlg1* deficient EAD clones expressing a dominant negative form of the *Drosophila* JNK Basket (*dlg1*^*G0342*^ *bsk*^*DN*^) (Fig 1H, 1I). In contrast, expression of a membrane tethered aPKC (aPKC^CAAX^) that causes mild overgrowth was sufficient to induce both Atf3::GFP and the ATRE reporter in specified wing disc compartments (*en>aPKC*^*CAAX*^ and *dpp>aPKC*^*CAAX*^) (Fig 1K, 1O). Although inhibiting JNK signaling suppressed the *aPKC*^*CAAX*^–mediated overgrowth, it did not prevent Atf3 induction (*en>aPKC*^*CAAX*^ *bsk*^*DN*^) (Fig 1L). Signaling via aPKC has been shown to engage Yorkie (Yki), a transcriptional co-activator in the Hippo pathway [10,25,28,29]. However, expression of a constitutively active form of Yorkie (*dpp>yki*^*act*^) did not induce the ATRE reporter (Fig 1P). Together, these findings highlight a role for Atf3 as a polarity response gene that is activated by loss of key polarity determinants downstream of aPKC but not JNK.

### Loss of *atf3* suppresses the effects of *dlg1* deficiency on eye development

Imaginal disc clones lacking either Scrib or Dlg1 surrounded by normal epithelium are severely disorganized, suffer from disturbed vesicular transport, lose the ability to terminally differentiate and are eliminated through cell competition involving JNK signaling [23,26,30]. While JNK and its downstream transcription factor Fos are required for apoptosis and suppression of Yki-mediated hyperproliferation of *scrib* and *dlg1* mutant cells [24,26,31-34], aPKC is responsible for the aberrant morphology and differentiation of the clonal cells [23].

Because Atf3 upregulation results from depletion of the Scrib complex components as well as from ectopic aPKC activation (Fig 1D, 1E, 1K), we investigated whether this excess Atf3 might contribute to the differentiation and morphological defects of EAD clones lacking Dlg1. While EADs bearing *dlg1* deficient clones (S1B, S1C Fig) produced significantly smaller, malformed adult eyes with patches of undifferentiated tissue (Fig 2A, 2C and S3A, S3F, S3G Fig), simultaneous removal of *atf3* (*atf3*^*76*^*dlg1*^*G0342*^) almost completely restored normal eye size and morphology (Fig 2D and S3B, S3F, S3G Fig), with only minor irregularities to the orderly hexagonal lattice (Fig 2D’). An equivalent genetic interaction was observed between *atf3* and the *dlg1*^*m52*^ allele [35,36] (S3D-G Fig). Importantly, adding a single copy of the *atf3*^*gBAC*^ transgene to animals with the double mutant mosaic EADs (*atf3*^*76*^*dlg1*^*G0342*^;; *atf3*^*gBAC*^/+) was sufficient to reinstate the aberrations to the adult eye morphology (S3C Fig), thus clearly showing that Atf3 is required for this *dlg1* deficiency phenotype to develop. Abnormalities were further exacerbated in adult eyes bearing *dlg1* deficient clones in which Atf3 was overexpressed (*dlg1*^*G0342*^ *atf3*^*wt*^) (Fig 2E). It is important to note that mosaic overexpression of Atf3 alone disturbed the normal ommatidial arrangement in the adult eye (Fig 2F). However, immunostaining of third instar EADs against a pan-neuronal marker Elav showed that, unlike cells lacking Dlg1, Atf3-expressing clones (*atf3*^*wt*^) differentiated (S4 Fig). In contrast, clonal loss of *atf3* alone (*atf3*^*76*^) did not markedly impact adult eye morphology (Fig 2B). These data provide causal evidence for the role of Atf3 downstream of disturbed epithelial polarity. They also indicate that while Atf3 is required for phenotypes caused by loss of *dlg1*, the gain of Atf3 alone does not fully recapitulate these defects.

**Fig 2.**
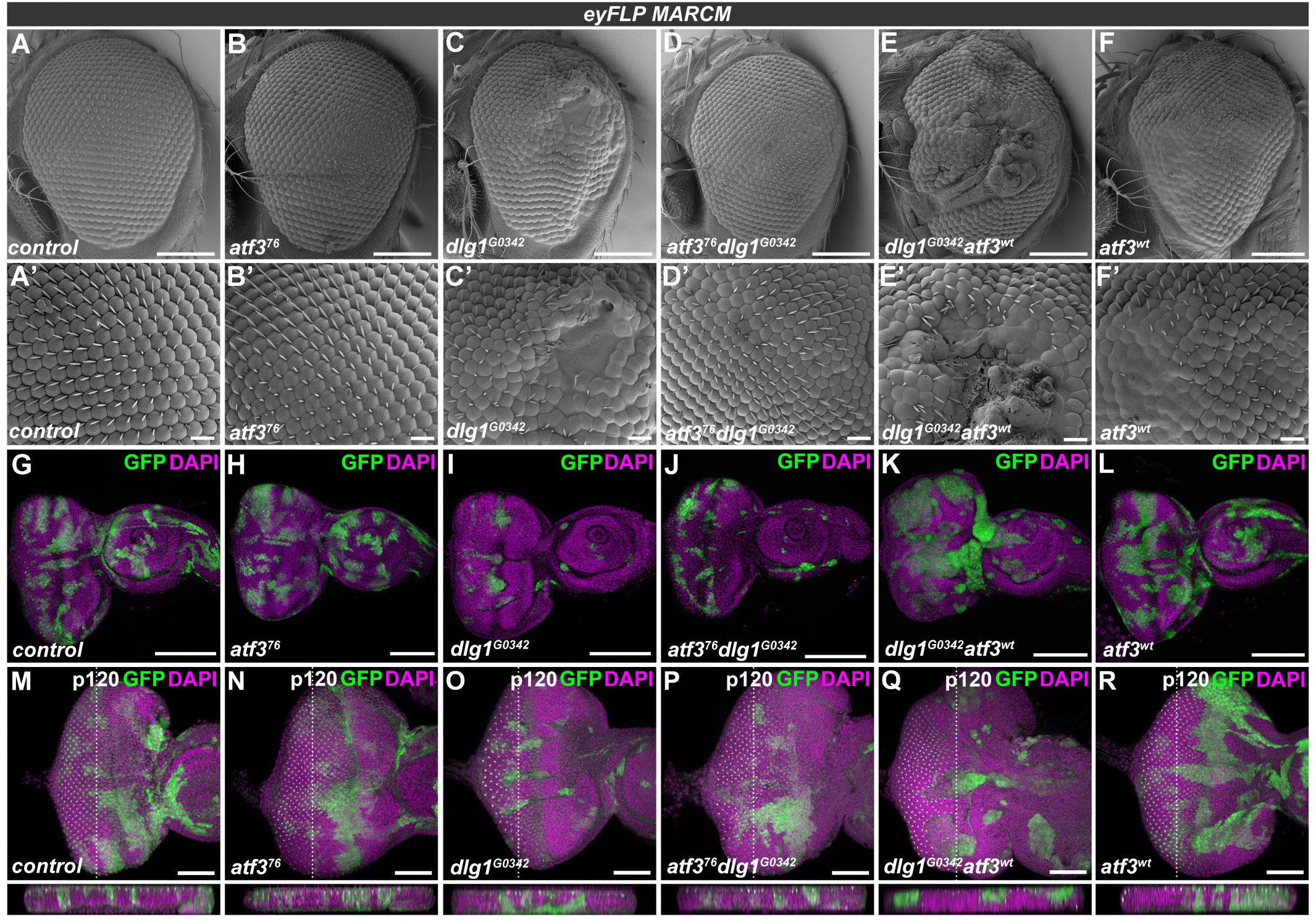
*atf3* genetically interacts with *dlg1*. **(A-R)** *eyFLP*-mediated mitotic recombination was used to generate clones (GFP) in EADs of the indicated genotypes. Lack of *dlg1* (C) produced smaller adult eyes containing patches of undifferentiated tissue (C’) relative to control eyes (A,A’) and those bearing *atf3*^*76*^ mutant clones (B,B’). Simultaneous clonal loss of *atf3* and *dlg1* (D) resulted in normally sized eyes with few to no differentiation defects (D’). In contrast, overexpression of *atf3* in *dlg1*^*G0342*^ clones (E) exacerbated the differentiation defects (E’). Clonal overexpression of *atf3* alone disturbed the orderly ommatidia array (F) generating glossy patches lacking bristles (F’). The *dlg1*^*G0342*^ (I) and *atf3*^*76*^*dlg1*^*G0342*^ (J) EAD clones were less abundant and smaller relative to control (G) and *atf3*^*76*^ mutant clones (H). Overexpression of *atf3* in *dlg1*^*G0342*^ cells generated clones with smooth boundaries composed of large, round cells (K). Overexpression of *atf3* alone did not impact clone shape or abundance (L). While no extrusion was observed either in case of *atf3*^*76*^ mutant (N) or *atf3* overexpressing (R) EAD clones compared to control EADs (M), the clonal loss of *dlg1* induced both apical and basal extrusion (O). Clones that lack *dlg1* and overexpress *atf3* are extruded apically (Q), while simultaneous loss of *dlg1* and *atf3* normalizes the epithelial integrity (P). EADs M-R were stained against the p120-catenin to aid the assessment of tissue architecture. Discs were counterstained with DAPI. Micrographs A-F are scanning electron micrographs. Micrographs G-R are projections of multiple confocal sections and show EADs 7 days after egg laying. Dotted lines indicate cross sections, which appear below the corresponding panels and are oriented apical side up. Scale bars: 100 μm (A-F), 10 μm (A’-F’), 100 μm (G-L), 50 μm (M-R).

### Loss of *atf3* restores polarity and differentiation to *dlg1* deficient cells independent of JNK activity

Closer examination of third instar larval EADs revealed the presence of fewer *dlg1*^*G0342*^ clones relative to control and *atf3*^*76*^ mosaic EADs (Fig 2G-I). Many of these mutant cells showed increased activity of the JNK-responsive TRE-DsRed reporter (S5B Fig). Interestingly, simultaneous loss of *atf3* neither restored the size or abundance of *dlg1*^*G0342*^ mutant cells nor prevented JNK activation (Fig 2J and S5A-C Fig). The exacerbated phenotype of *dlg1*^*G0342*^*atf3*^*wt*^ mosaic adult eyes correlated with the presence of contiguous clonal patches in third instar EAD that were markedly rounder compared to clones lacking *dlg1* alone (Fig 2I, 2K). Although the individual clones appeared larger, the overall amount of GFP positive tissue was not increased when compared to *dlg1* mutant clones (S5A Fig). The cross sections of the eye primordia further revealed that in contrast to *dlg1* deficient cells which accumulated basally, the majority of *dlg1*^*G0342*^*atf3*^*76*^ clones remained in the disc proper similar to control and *atf3* mutant cells (Fig 2M-P). Strikingly, overexpression of Atf3 in cells lacking *dlg1* promoted their apical extrusion (Fig 2Q). To rule out that clone elimination contributed to the observed rescue phenotype in the *atf3*^*76*^*dlg1*^*G0342*^ adult eyes, we took two alternative approaches. To inhibit death of clonal cells, we expressed the baculoviral caspase inhibitor p35 in *dlg1*^*G0342*^ and *atf3*^*76*^*dlg1*^*G0342*^ EAD clones. To reduce competition elicited by neighboring cells, we utilized the EGUF/hid technique [37], which facilitates expression of a pro-apoptotic protein Hid in the non-clonal cells within the Glass multiple reporter (GMR) domain causing their elimination during pupal stages [38]. As expected, blocking apoptosis by p35 increased the abundance of mutant clonal cells to a similar extent in both genotypes (S5A Fig). Importantly, nearly four times more adults bearing *atf3*^*76*^*dlg1*^*G0342*^*p35* mosaic EADs emerged compared to *dlg1*^*G0342*^*p35* animals (S5D Fig). Adult eyes derived from *dlg1*^*G0342*^*p35* EAD contained only small patches of differentiated photoreceptors compared to controls (S5E, S5F Fig), whereas *atf3*^*76*^*dlg1*^*G0342*^*p35* eyes were either similar to those of *atf3*^*76*^*dlg1*^*G0342*^ animals (Fig 2D and S5G Fig) or exhibited modest differentiation defects (S5H Fig). Consistently, the overall morphology and differentiation pattern was markedly improved in adult eyes composed entirely of *atf3*^*76*^*dlg1*^*G0342*^ mutant tissue generated by the EGUF/hid method. In contrast to only remnants of differentiated tissue in *dlg1*^*G0342*^ mutant eyes, clear ommatidial arrays were observed in control and *atf3*^*76*^*dlg1*^*G0342*^ adult eyes, (S5I-K Fig). Taken together, these data strongly argue against apoptosis being the basis of the genetic rescue conferred by *atf3* deficiency in *dlg1* mutant clones, and instead suggest that Atf3 contributes to cell extrusion and differentiation defects caused by loss of Dlg1.

To better characterize how loss of *atf3* impacts *dlg1* mutant phenotypes, we stained third instar larval EADs for markers of differentiation and cell adhesion. Unlike control clones, *dlg1*^*G0342*^ mutant cells located posterior to the morphogenetic furrow of the eye primordium frequently lacked the pan-neuronal marker Elav. Many of them flattened and delaminated, pushing some of the non-clonal Elav-positive cells to the basal side of the epithelium (Fig 3A, 3B). In contrast, *atf3*^*76*^*dlg1*^*G0342*^ clones remain columnar, contributing to photoreceptor and interommatidial cell differentiation (Fig 3C). Importantly, immunostaining for the apical determinant Crumbs, the adherens junction protein DE-cadherin (DE-cad) and the lateral membrane marker Fasciclin III (FasIII) revealed a clear rescue of the apicobasal organization of the ommatidia in *atf3*^*76*^*dlg1*^*G0342*^ clones compared to the disturbed architecture in clones mutant for *dlg1*^*G0342*^ alone (Fig 3A-F). These findings uncover novel genetic interaction between Atf3 and the central Scrib polarity module component Dlg1 and establish Atf3 as a driver of the structural and differentiation defects stemming from the loss of *dlg1*.

**Fig 3.**
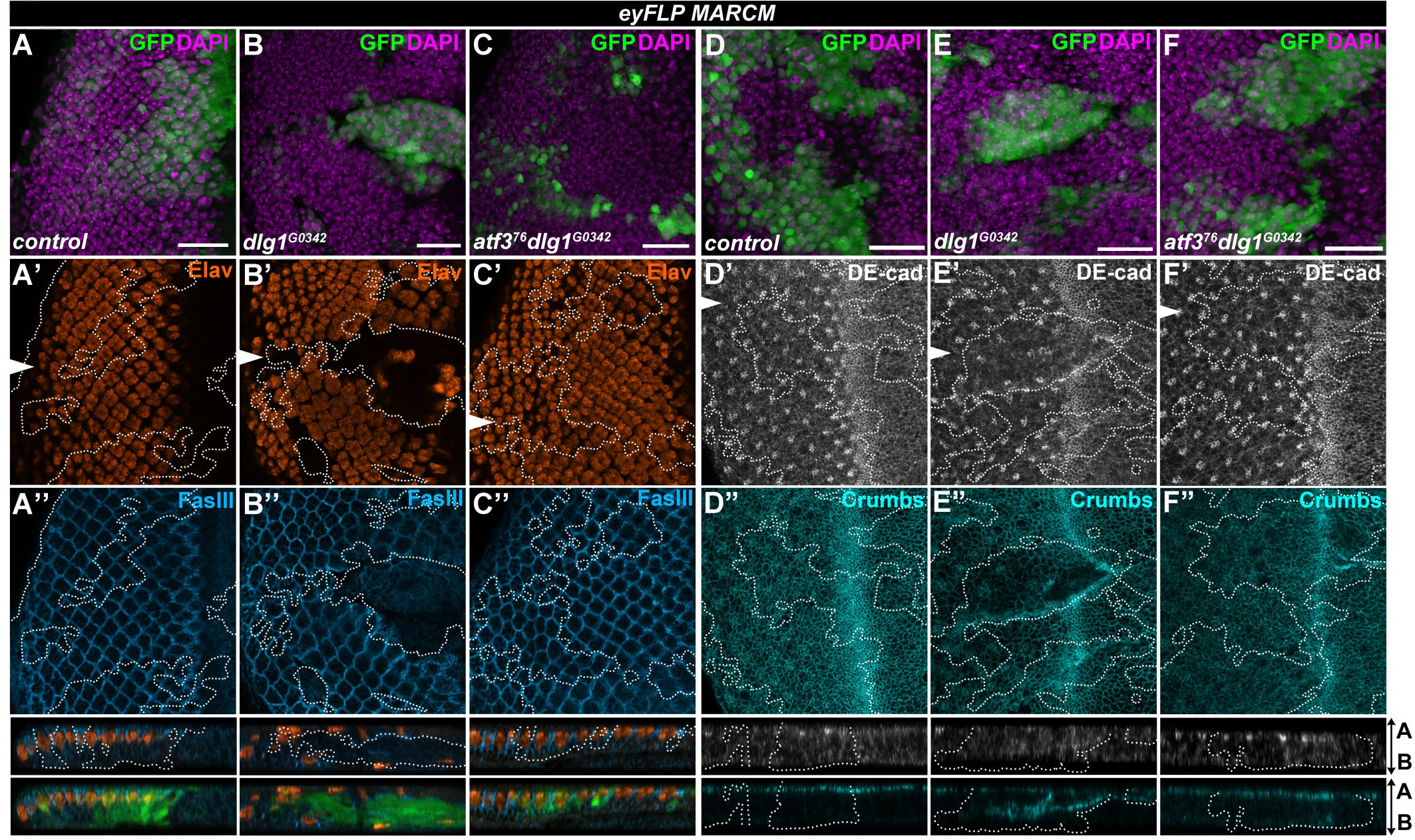
Loss of *atf3* restores differentiation and polarity to *dlg1* mutant clones. **(A-F)** *eyFLP*-mediated mitotic recombination was used to generate clones (GFP) in the EADs of the indicated genotypes. **(A-C)** Immunostaining shows the stereotypical pattern of differentiated photoreceptors (Elav) and the lateral membrane component Fasciclin III (FasIII) in control discs (A’-A’’). Patterns of these markers are disturbed in *dlg1*^*G0342*^ clones (B’- B’’), and restored in *atf3*^*76*^*dlg1*^*G0342*^ clones (C’- C’’). Similar to control (A’’), *atf3*^*76*^*dlg1*^*G0342*^ cells (C’’) show columnar organization and maintain contact with the apical surface while *dlg1*^*G0342*^ clones delaminate (B’’). **(D-F)** The normal patterns of the adherens junction component DE-cadherin (DE-cad) (D’) and the apical determinant Crumbs (D’’) are disturbed in *dlg1*^*G0342*^ clones (E’-E’’) but normalized in *atf3*^*76*^*dlg1*^*G0342*^ clones (F’-F’’). Discs were counterstained with DAPI. White arrows indicate cross sections, which appear below the corresponding panels and are oriented apical side up. Clones are outlined with white dotted lines. Micrographs are otherwise single confocal slices. All images show EADs 7 days after egg laying. Scale bars: 20 μm (A-F).

### Disturbances to the endosomal trafficking machinery in *dlg1* mutant epithelia require Atf3

Besides altered differentiation and mislocalization of polarity proteins, disturbed cellular trafficking [30,39-41] is a hallmark feature of epithelial cells lacking the Scrib module components. Therefore, we tested if the distress of the trafficking machinery upon *dlg1* loss was Atf3-dependent. Interestingly, neither *dlg1*^*G0342*^ nor *atf3*^*76*^*dlg1*^*G0342*^ EAD clones enlarged by p35 expression showed changes in the uptake of fluorescently labeled dextran compared to surrounding tissue (S6A-C Fig), indicating that endocytic activity in cells lacking *dlg1* was not affected. However, the staining of intracellular vesicle components revealed marked changes to the amount and distribution of early and recycling endosomes. While levels of the Rab5-positive vesicles were reduced in *dlg1*^*G0342*^ and *dlg1*^*G0342*^*p35* mutant cells located in front or behind the morphogenetic furrow (Fig 4A’, 4B’ and S7B, S7E Fig), the pool of Rab11 recycling endosomes increased (Fig 4A”, 4B”). In line with these findings, Dlg1-deficient cells also displayed an abnormal distribution of the Syntaxin7/Avalanche (Avl)-positive vesicles (S6E Fig). In contrast, the Rab5, Avl and Rab11 staining patterns in *atf3*^*76*^*dlg1*^*G0342*^ double mutant EAD clones were comparable to those seen in control clones or surrounding normal tissue (Fig 4C, 4D and S6D, S6F, S7A, S7C, S7D, S7F Fig). Strikingly, the changes to the endosomal compartment observed in *dlg1* mutant tissue could be recapitulated by the Atf3 overexpression. While apical levels of Rab5 were decreased, Rab11-marked vesicles were enriched on the basal side of the wing imaginal discs in cells expressing Atf3 when compared to surrounding tissue (S8A, S8B Fig). In addition, immunostaining of wing discs bearing Atf3-expressing clones revealed changes in levels and localization of polarity proteins whose proper membrane placement requires a functional trafficking machinery [42,43]. While the apical levels of Crumbs were markedly lower, the integrin subunit Myospheroid (Mys), which normally is restricted to the basal cell surface, was detected along the entire lateral membrane in Atf3-expressing clones relative to surrounding tissue (S8C, S8D Fig). These data demonstrate that Atf3 contributes to the alterations of the endosomal machinery upon loss of polarity and its overexpression is sufficient to mimic some of the hallmark features of *dlg1*-deficient epithelial cells. Interestingly, despite the shift in apicobasal polarity markers Atf3-expressing cells, unlike those lacking *dlg1*, maintained their columnar shape and were not extruded from the epithelium (Fig 2R and S8 Fig).

**Fig 4.**
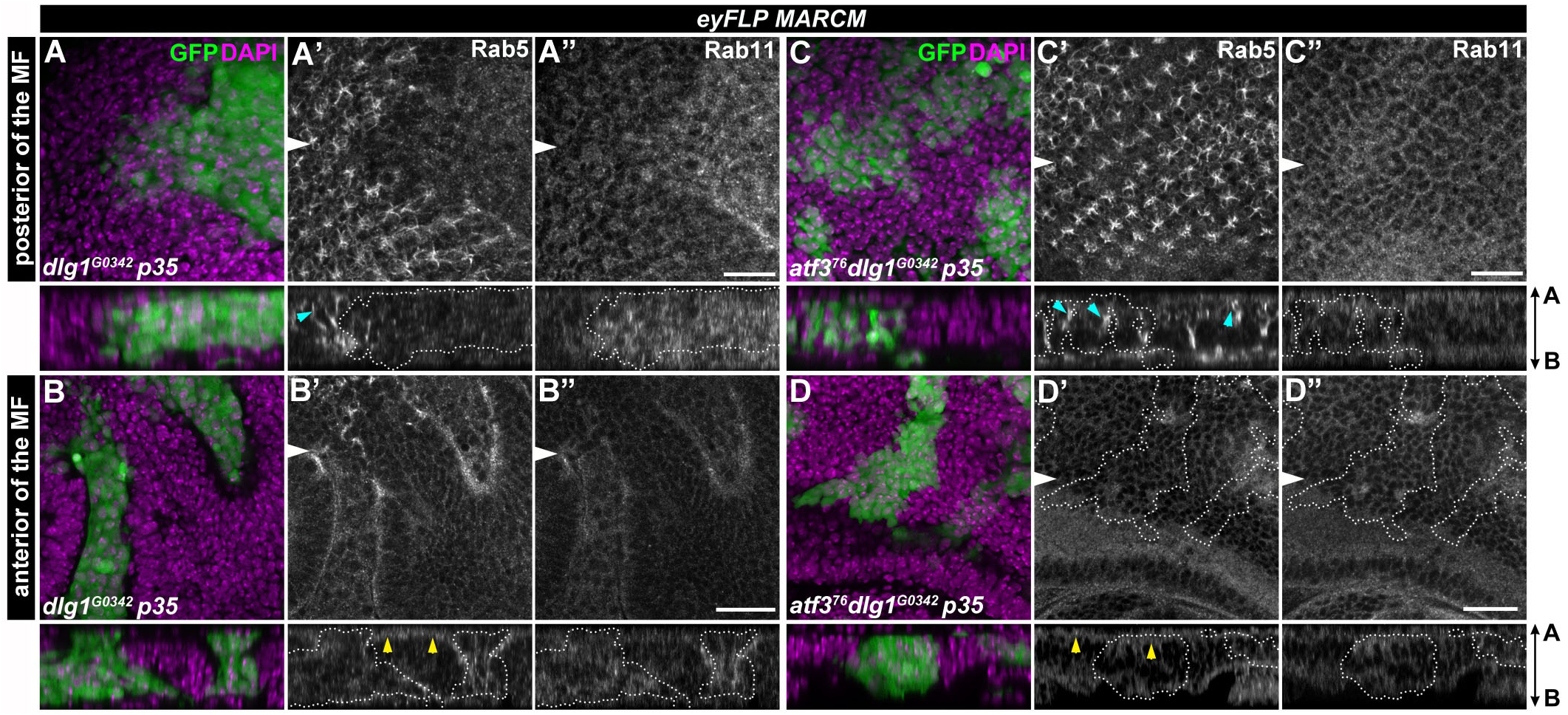
Impairment of Rab5 and Rab11 abundance and localization in *dlg1* mutant clones is Atf3-dependent. **(A-D)** *eyFLP*-mediated mitotic recombination combined with a co-expression of p35 was used to generate larger clones (GFP) in the EADs of the indicated genotypes. The levels and localization of early Rab5-positive endosomes are disturbed upon loss of *dlg1* both posterior (A’) and anterior (B’) to the morphogenetic furrow, while recycling Rab11 vesicles are enriched in *dlg1* mutant clones relative to surrounding tissue (A’’,B’’). Both Rab5 and Rab11 vesicle amounts and distribution are normalized in *atf3*^*76*^*dlg1*^*G0342*^ double mutant clones (C-D). On the cross sections, blue arrowheads indicate the regular Rab5 enrichment along the photoreceptors (A’,C’), yellow arrowheads indicate the apical localization of Rab5 in epithelial cells anterior of the MF (B’,D’). Note that both patterns are lost in *dlg1* deficient cells. Discs were counterstained with DAPI. White arrows indicate cross sections, which appear below the corresponding panels and are oriented apical side up. Clones are outlined by white dotted lines. All images show EAD 7 days after egg laying. Scale bars: 10 μm (A,C), 20 μm (B,D).

### Atf3 binds a 12-nucleotide motif in genes fundamental to cytoarchitecture

As a bZIP transcription factor, Atf3 is expected to regulate gene expression through binding to specific DNA sequences. To capture a snapshot of genomic regions bound by Atf3, we employed the Atf3::GFP fusion protein expressed from the recombineered *atf3*^*gBAC*^ to perform chromatin immunoprecipitation followed by high throughput sequencing (ChIP-seq). The ChIP-seq using an anti-GFP antibody identified 152 genomic locations significantly enriched in samples prepared from *atf3*^*76*^;; *atf3*^*gBAC*^/+ mated adult males compared to *yw* controls (Fig 5A and S1 Dataset). Using the Multiple Em for Motif Elicitation (MEME) tool [44], we derived a 12-column position weight matrix (PWM) yielding a palindromic binding site which we call the *Drosophila* Atf3 motif (Fig 5B and S2 Dataset). This motif occurred in 112 of the 152 significantly enriched peaks in Atf3 samples, which correspond to 121 genes whose cis-regulatory regions contained an Atf3 motif (Fig 5A and S1 Dataset). Submission of this PWM to Tool for Motif to Motif comparison (TOMTOM) [45] determined that Atf3 recognizes DNA that is most similar to the Atf2 binding motif in *Drosophila* and ATF3/JDP2 motif in humans (Fig 5B). Closer examination of the Atf3-response elements revealed an 8-nucleotide core that closely resembles the cyclic AMP response element (CRE) (TGACGTCA) (Fig 5B). We have previously reported that the bZIP domain of the *Drosophila* Atf3 protein indeed directly binds this DNA element [6]. Our present experiments using the ATRE and mATRE reporters further confirm that Atf3 can control expression from this site in cultured cells and *in vivo* (Fig 1M’ and S2 Fig).

**Fig 5.**
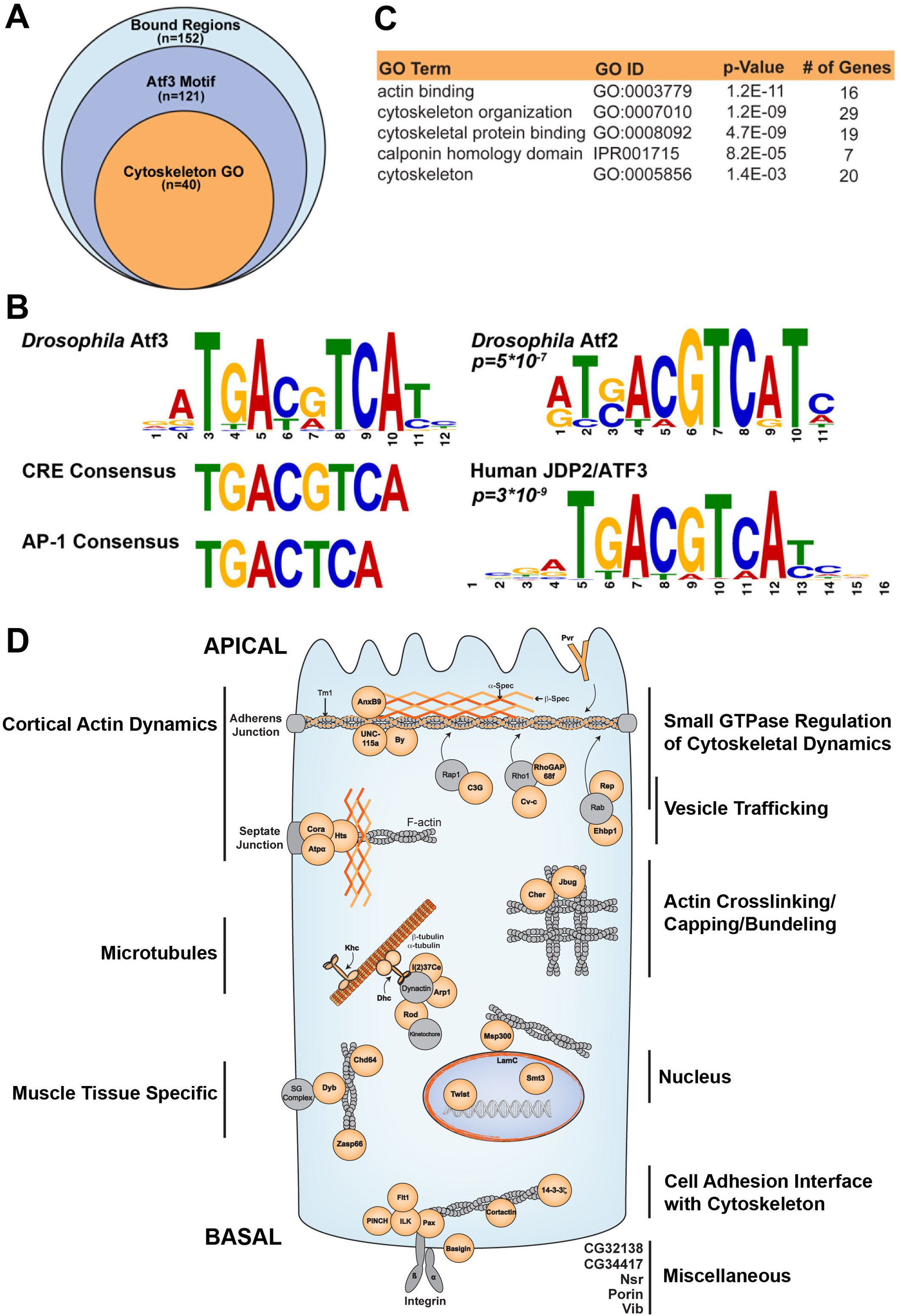
Atf3 targets genes fundamental to epithelial cytoarchitecture. **(A)** ChIP-seq identified 152 genomic positions bound by Atf3 in samples from *atf3*^*76*^/*Y;;atf3*^*gBAC*^/+ adult flies. **(B)** 112 genomic regions, proximal to 121 genes, contained the *Drosophila* Atf3 motif. The core of the Atf3 motif closely matches the CRE element but not the AP-1 consensus site. The Atf3 motif most closely resembles the *Drosophila* Atf2 motif (*p=5E-7*) and human JDP2/ATF3 motif (*p=3E-9*). **(C)** Of the 121 genes targeted by Atf3, 40 have roles in cytoskeleton organization and dynamics. **(D)** A schematic shows Atf3 targets (orange) belonging to the cytoskeleton GO cluster broken into subgroups and placed within representative spatial contexts in an epithelial cell.

The gene ontology (GO) analysis with FlyMine [46] revealed that structural and regulatory components of the cytoskeleton were significantly enriched among Atf3-bound regions. In total, 40 of the identified 121 genes containing Atf3 motifs belong to at least one out of five GO terms corresponding to cytoskeleton regulation (Fig 5A, 5C and S1 Dataset). Figure 5D depicts a summary of putative Atf3 targets engaged in their corresponding functions in a model epithelial cell. Importantly, an independent Atf3 ChIP experiment from third instar larval EADs determined that several of these targets including microtubule subunits (*alphaTub84B* and *betaTub56D*) and nuclear *Lamin C* (*LamC*) were also bound by the Atf3 in imaginal epithelia (S9A, S9B Fig) suggesting a potential mechanistic basis for the Atf3 driven phenotypes in epithelial cells with compromised apicobasal axis.

### Atf3 overexpression has a broad effect on gene transcription in imaginal disc epithelia

To complement the ChIP-seq approach and characterize the transcriptional response to excess Atf3, we compared the transcriptomes of mosaic EADs overexpressing Atf3 with control expressing only GFP. Strikingly, RNA-seq revealed a widespread shift in the expression of 4957 genes (S3 Dataset), with nearly three times as many transcripts being downregulated (n = 3666) than enriched (n = 1291) by fold change ≥1.5 relative to control (Fig 6A and S3 Dataset).

**Fig 6.**
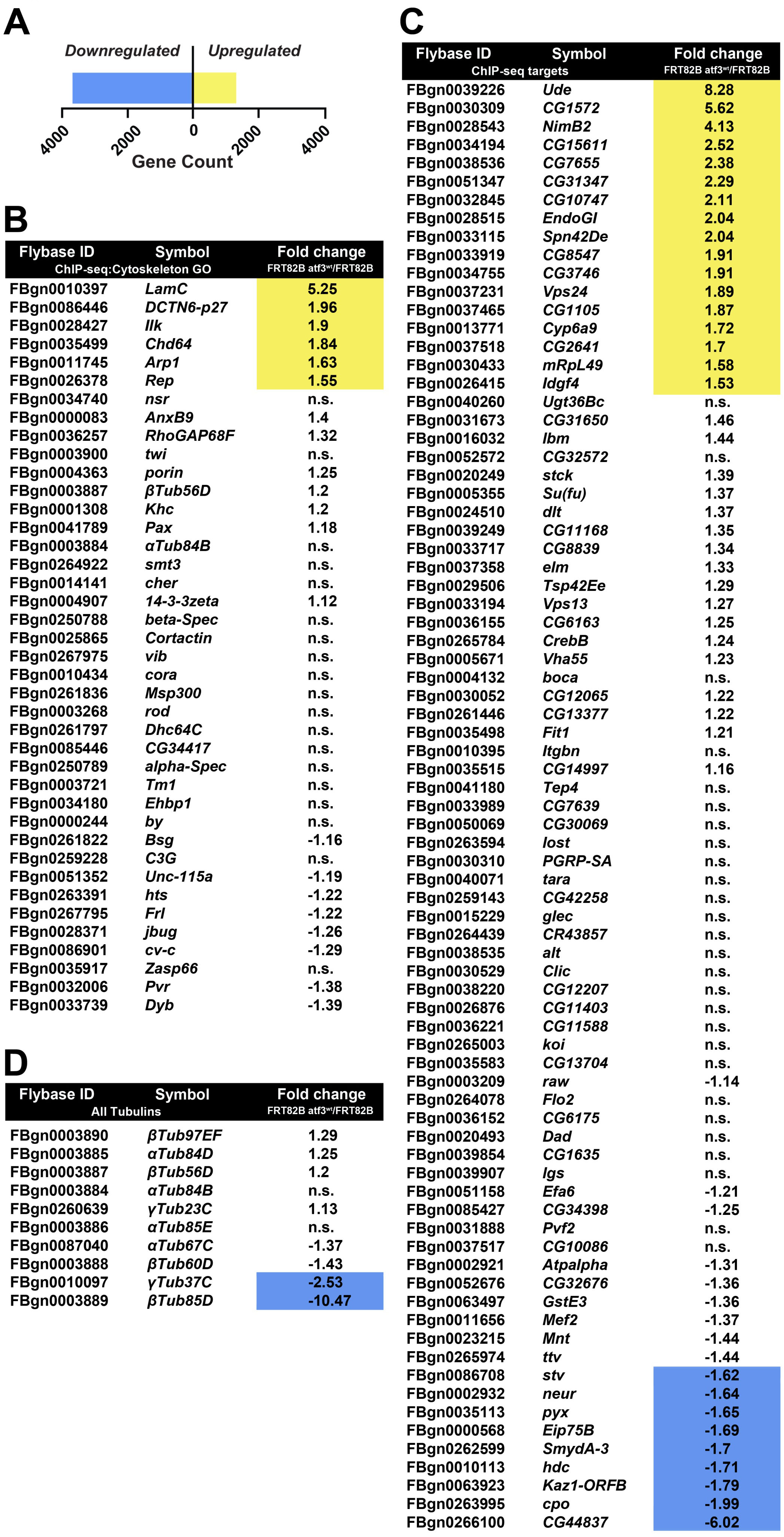
Gain of Atf3 induces a broad transcriptional response in EAD epithelium. **(A)** Transcriptome analysis identified a total of 4957 genes that were differentially regulated by a factor ≥1.5 in mosaic EADs expressing Atf3 (*eyFLP*^*MARCM*^ *atf3*^*wt*^). Three times as many genes were transcriptionally repressed (n=3666) than activated (n=1291). **(B-C)** Tables list the Flybase ID, gene symbol (Symbol), and relative fold change in gene expression (Fold Change *FRT82B atf3*^*wt*^/*FRT82B*) of the Atf3 targets identified by ChIP-seq that are part of the cytoskeleton GO cluster (B) and the 80 Atf3 ChIP-seq targets which do not belong to the cytoskeleton GO cluster (C). **(D)** A list summarizing the regulation of all tubulin genes in mosaic EADs overexpressing Atf3 sorted in descending order of transcript abundance.

Of the 121 genes identified as Atf3 targets by ChIP-seq in the adults, 32 were differentially expressed at the mRNA level (Fig 6B and 6C) in *atf3*^*wt*^ mosaic EADs, most of which (23) showed transcriptional upregulation including six genes (*Arp1, Chd64, DCTN6-p27, Ilk, LamC, Rep*) that belonged to the cytoskeleton GO cluster (Fig 6B). In accordance with the mRNA-seq, an independent qRT-PCR of selected Atf3 ChIP-seq targets detected enrichment of *LamC, ude, DCTN6-p27* and *Arp1* transcripts while *alphaTub84B* and *betaTub56D* remained unchanged (S9A Fig) despite being occupied by Atf3 in both the adults and EADs (S1 Dataset and S9B Fig). To identify additional putative Atf3 targets among genes misregulated in the RNA-seq dataset, we scanned the *Drosophila* genome with our experimentally derived Atf3 PWM using the Find Individual Motif Occurrences (FIMO) [47] and Peak Annotation and Visualization (PAVIS) [48] utilities. The Atf3 motif (Fig 5B) was found within 5 kb upstream and 1 kb downstream of 1252 genes differentially regulated in *atf3*^*wt*^ mosaic EADs (S3 Dataset). Importantly, a significant proportion of transcripts altered by Atf3 expression in the mosaic EADs overlapped with genes that had been found deregulated in either *scrib* or *dlg1* mutant wing imaginal discs [10] (S10 Fig and S3 Dataset). This intersection shows that the gain of Atf3 and the loss of the Scrib polarity module elicited a partly overlapping genetic response.

### Atf3 controls the microtubule network and Lamin C downstream of loss of polarity

The requirement of Atf3 for the manifestation of cytoarchitecture and polarity defects in *dlg1* deficient cells prompted us to further focus on its regulation of microtubules and LamC, the primary building blocks of the cytoskeleton and nucleoskeleton, respectively. Microtubules are classically associated with controlling cell shape and division, but they are also central to polarity by serving as railroad tracks for directed vesicular trafficking [49]. Nuclear lamins on the other hand provide structure and stiffness to the nuclear envelope [50,51]. The coupling between the cytoskeleton and the nuclear lamina via the LINC-complex is essential for maintaining the mechanical properties of the cell and signal transduction [52]. Immunostaining for LamC showed that the increase in *LamC* transcription in response to excess Atf3 (Fig 6B and S9A Fig and S3 Dataset) translated into a dramatic enrichment of the LamC protein in the nuclear envelope of EAD or wing disc cells overexpressing Atf3 (Fig 7A, 7D). Importantly, LamC protein levels also increased in the wing disc epithelium deficient for Scrib (*en>scrib*^*RNAi*^) and in *dlg1* mutant clones of the EAD (Fig 7B, 7E), whereas they were markedly reduced in *atf3*^*76*^*dlg1*^*G0342*^ double mutant EAD clones compared to the surrounding tissue and control clones (Fig 7C, 7F). In contrast to *LamC* enrichment, expression of at least two microtubule genes (*βTub85D* and *γTub37C*) was downregulated in *atf3*^*wt*^ mosaic EADs (Fig 6D, S3 Dataset). Interestingly, immunostaining for α- and β-Tubulin revealed a reduced microtubule network and markedly lowered levels of non-centrosomal γ-Tubulin in Atf3-overexpressing clones of the wing imaginal disc relative to surrounding epithelial tissue (Fig 8A-C). In addition, eye primordia bearing *dlg1* mutant clones showed abnormal microtubule organization that was partially normalized by simultaneous clonal loss of *atf3* (Fig 8D-F). Taken together, our molecular and genetic approaches identified microtubule encoding genes and LamC as new targets of Atf3. We demonstrate that the upregulation of LamC and disturbances to microtubule network are the common hallmarks of epithelial cells in which Atf3 activity is enhanced either by transgenic overexpression or as a result of disturbed polarity. While LamC is a *bona fide* target of Atf3, changes to microtubule network likely arise indirectly as a part of the secondary response to ectopic Atf3 activity.

**Fig 7.**
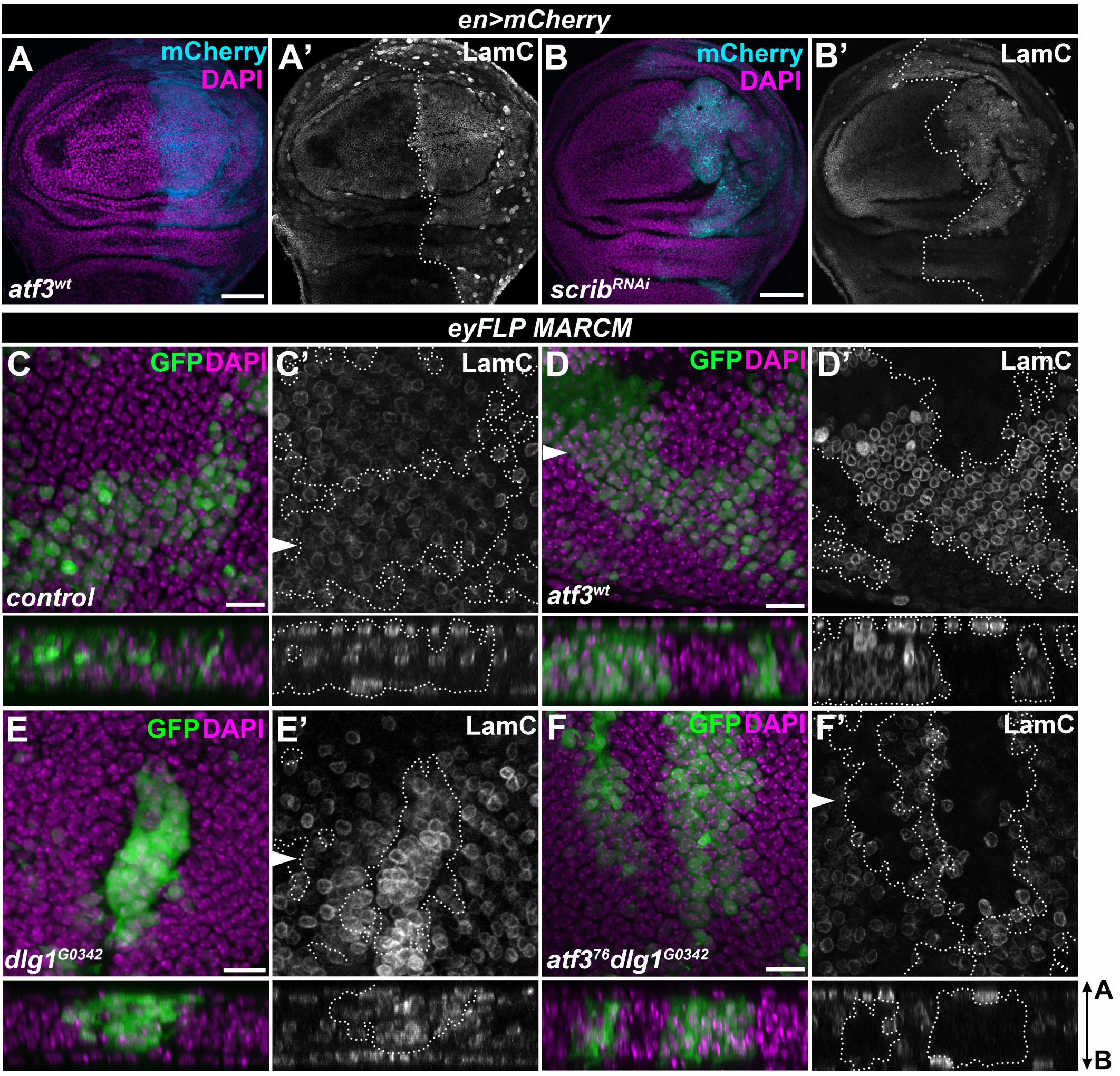
LamC is a *bona fide* target of Atf3. **(A-B)** Both Atf3 overexpression (*en>atf3*^*wt*^) and Scrib knockdown (*en>scrib*^*RNAi*^) increased LamC levels in the posterior compartment of the wing imaginal disc (mCherry) (A’,B’). **(C-D)** *eyFLP*-mediated clonal overexpression of Atf3 (GFP) markedly upregulated the LamC protein (D’) compared to levels detected in surrounding tissue and control clones (C’). **(E-F)** LamC was also enriched in *dlg1*^*G0342*^ clones (E’), but was reduced below control levels in *atf3*^*76*^*dlg1*^*G0342*^ cells (F’). Discs were counterstained with DAPI. Images represent single slices from confocal stacks. The wing disc posterior compartments (A-B) and EAD clones (C-F) are outlined with white dotted lines. On images C’-F’, the arrows indicate cross sections shown below the corresponding images, with the apical side facing up. Scale bars: 50 μm (A-B), 10 μm (C-F).

**Fig 8.**
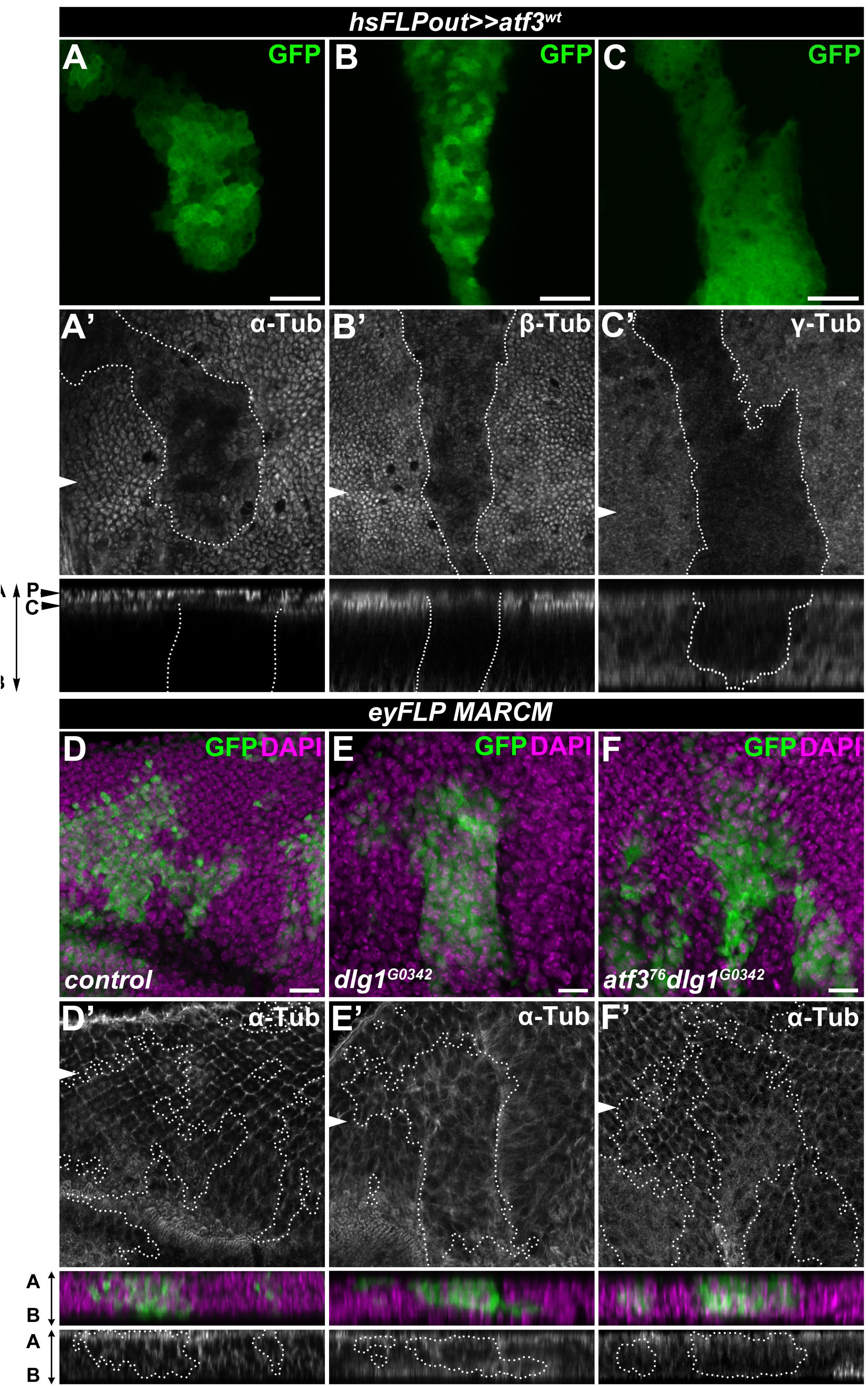
Increased Atf3 levels disturb the microtubule network. **(A-B)** Columnar cells in the wing disc expressing Atf3 (*hsFLPout>>atf3*^*wt*^) (GFP) contained less apical, subcortical α- and β-Tubulin compared to surrounding tissue (A’,B’) and non-clonal cells of the peripodial epithelium (P) directly above (A’). **(C)** Wing disc clones expressing Atf3 display lower levels of non-centrosomal γ-Tubulin (C’). **(D-F)** *eyFLP*-mediated mitotic recombination was used to generate clones (GFP) in the EADs of the indicated genotypes. Clones lacking *dlg1* (E) show disturbed α-Tubulin localization (E’) compared to control clones (D-D’) in the photoreceptor region of the EADs. Clones lacking both *dlg1* and *atf3* (F) show normalized microtubule distribution along the apicobasal axis (F’). Discs were counterstained with DAPI. The arrows indicate cross sections shown below the corresponding images, with the apical side facing up. Clones are outlined with white dotted lines. Scale bars: 10 μm (A-F).

## Discussion

Classical insults to epithelial polarization include disruptions of the Crumbs, aPKC/Par, or Scrib protein modules, which all alter tissue organization, proliferation, differentiation, and cell viability [1]. The engagement of transcription factors presents a significant but poorly understood consequence to compromised polarity [10,23,28,29,53,54]. In this study, we identify Atf3 as a novel polarity response gene induced by aPKC signaling upon loss of the neoplastic tumor suppressors of the Scrib polarity module. Our results exclude JNK activation stemming from loss of polarity as a driver of *atf3* expression. Blocking JNK in *dlg1* mutant clones or tissues with active aPKC signaling did not abrogate Atf3 expression. Furthermore, activation of JNK signaling neither induced Atf3 in the wing imaginal disc, nor significantly activated the ATRE reporter in S2 cells. Moreover, activated Yki, a downstream effector of aPKC [10,28,29] and a context-dependent mediator of JNK signaling [24,25,54] was not sufficient to induce Atf3 activity. Future studies should determine which transcription factors downstream of aPKC regulate *atf3* expression.

This study positions Atf3 as a major regulator in the aPKC circuit, as removing Atf3 from *dlg1* deficient cells recapitulated the phenotypes conferred by reduced aPKC activity in *dlg1* tissue, including restoration of a polarized, columnar morphology and differentiation in the eye primordium [23]. Importantly, Atf3 acts in a manner distinct from that of the bZIP protein Fos and the TEAD/TEF transcription factor Scalloped (Sd) and its co-activator Yki, operating downstream of disrupted polarity [23,28,29,32,54]. In *scrib* and *dlg1* mutant cells, Fos and Sd/Yki promote apoptosis and proliferation, respectively; however they are not linked to disturbed cytoarchitecture [24,28,33]. Conversely, lack of Atf3 does not block JNK activity or the elimination of *scrib/dlg1* mutant clones, whereas overexpression of dominant-negative forms of JNK or removal of Fos do [23,26,32]. As loss of *atf3* did not affect *dlg1* clonal abundance, we conclude that *atf3* deficiency neither reduces the proliferative potential of *dlg1* tissue nor interferes with JNK-mediated repression of Yki [24]. Taken together, we propose that Atf3, Fos, and Sd/Yki act downstream of cell polarity insults in parallel (Fig 9).

**Fig 9.**
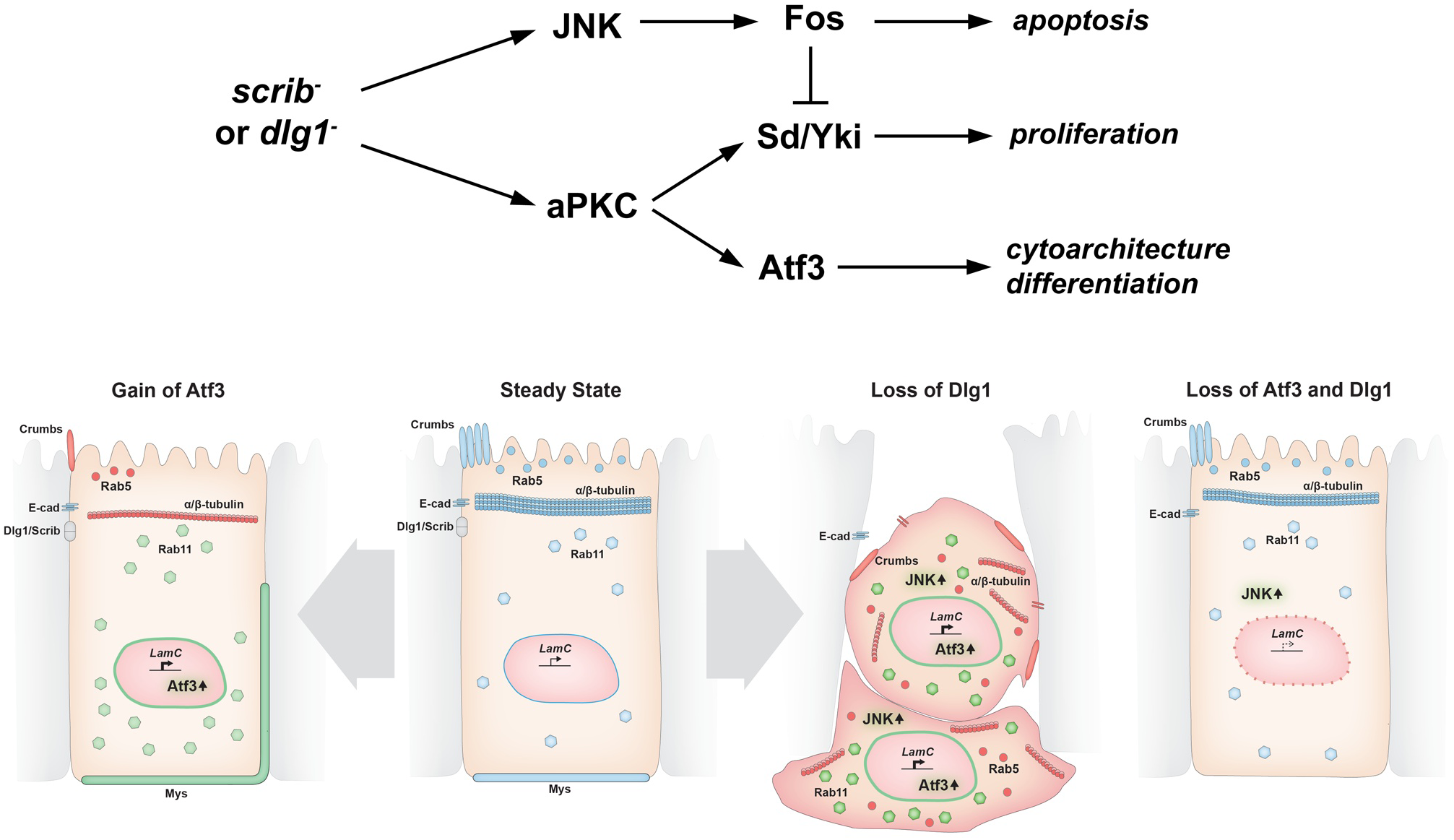
Model depicting the effects of Atf3 misregulation in epithelia. In our working model (upper diagram), loss of *dlg1/scrib* activates aPKC and JNK signaling. Through Fos, JNK induces apoptosis and restrains excessive cell proliferation that would result from aPKC-mediated activation of Yki. Dlg1 mutant cells flatten and are extruded. Aberrant aPKC activity induces Atf3 expression independent of JNK, leading to defects in cell morphology and differentiation. Columnar cell organization and the aberrant distribution of Rab5 and Rab11 vesicles and polarity proteins observed upon loss of Dlg1 are rescued by simultaneous loss of Atf3, while JNK signaling remains active. Atf3 overexpression negatively impacts the microtubule network, dysregulates Rab5 and Rab11 trafficking routes, and perturbs the distribution of apicobasal polarity proteins. Nevertheless, Atf3 overexpressing cells remain columnar. Note that in all cases, LamC is regulated in an Atf3-dependent manner. Physiological levels/correct localization are indicated in blue, increased levels/expanded localization in green and reduced levels/mislocalization in red.

Impairment of the vesicular trafficking machinery has been recognized as a key feature of epithelial cells lacking polarity determinants such as Crumbs or the components of the Scrib complex [30,39,41,55]. However, the deregulation of endocytosis has been also shown to drive polarity defects and to alter signal transduction resulting in overgrowth or apoptosis depending on the tissue and genetic context [30,39,41,56]. Here, we show that while bulk endocytosis is not affected in *dlg1* mutant clonal cells of the EAD, levels and distribution of specific components of the vesicle trafficking machinery and their cargo are disturbed and these defects depend on Atf3. However, despite the clear rescue of the Rab5 and Rab11 trafficking apparatus in *atf3 dlg1* double mutant clones JNK signaling remains upregulated and cells are eliminated, suggesting that multiple mechanisms are in play to promote JNK activity upon loss of *dlg1*. Importantly, the disruption of Rab5 and Rab11-mediated trafficking routes can be recapitulated by Atf3 overexpression. The reduced levels of Crumbs, whose membrane delivery requires vesicular transport from the *trans* Golgi-network [30], and the ectopic expansion of Mys (βPS-integrin) along the lateral membrane, whose turnover relies on Rab5 and Rab21 vesicles [57], further highlight the impact of enhanced Atf3 activity on the trafficking and recycling of cellular and membrane components. Whether this contribution is direct or indirect via regulating cellular architecture remains to be determined.

Our study provides initial *in vivo* insights into the binding specificity of *Drosophila* Atf3. The ChIP-seq experiment revealed an enrichment of Atf3 targets among the constituents of the cytoskeleton and factors that are associated to it. Atf3 bound and deregulated the expression of several genes encoding tubulin subunits and tubulin-associated motor proteins (ChIP-seq: *αTub84B, βTub56D, Khc, Dhc64C*; mRNA-seq: *βTub85D* and *γTub37C*). Importantly, the microtubules and motors are not only pivotal to the maintenance of cellular structure, but also to intracellular trafficking processes [49,58,59]. Atf3 upregulation, whether resulting from loss of apicobasal polarity or induced by transgenic overexpression, disrupts the stereotypical tubulin organization and impairs vesicle trafficking. This suggests that Atf3 is crucial for proper cytoarchitecture and the associated transport mechanisms in epithelial cells.

In addition, Atf3 targeted genes responsible for the organization and dynamics of the actin cytoskeleton (*cher, jbug, cortactin*), as well as genes linked to the integrin adhesion complex (*Itgbn, Ilk, Stck, Fit1, Pax*). The dynamic integrin-actin coupling is essential for sensing and transducing mechanical stimuli, which result in the adjustment of the intracellular tension and changes in cell behavior and morphology. On the other hand, altered mechanotransduction has been linked to polarity defects and tumorigenesis [60-63]. Interestingly, we show that the extrusion process which relies on coordinated activities of the cytoskeleton and adhesion modules can be modulated by Atf3 gain or loss. Thus, it is plausible that cellular shape and tension are controlled by Atf3 in the context of disturbed polarity.

Finally, we identified Atf3 targets among the components of the nuclear lamina (*LamC*), and the nuclear-envelope-spanning linker of nucleoskeleton and cytoskeleton (LINC) complex (*Msp300, koi*). While a filamentous meshwork of nuclear lamins underlies the inner nuclear membrane and provides structural properties to the nuclear envelope [50,51] the LINC complex couples the nucleoskeleton to both the actin and microtubule networks [52]. Importantly, the LINC complex and nuclear lamins have emerged as central components of the mechanosensing and mechanotransduction machinery, not only relaying mechanical forces to the nucleus [64], but also acting as force generators, ultimately inducing cytoskeletal rearrangement, polarization and cell shape changes [65].

With RNA-seq profiling we aimed to defined a gene expression program provoked by increased Atf3 levels in primordial epithelia. Interestingly, transcriptome analysis revealed that only a quarter of Atf3 ChIP-seq targets were misexpressed in *atf3*^*wt*^ mosaic EADs, which represent a fraction of all differentially regulated genes. This relatively small intersection between ChIP-seq targets identified in adult flies and the EAD transcriptome could reflect a tissue specific context for DNA-binding and transcriptional regulation by Atf3. Moreover, the widespread transcriptional deregulation following Atf3 overexpression in epithelia indicates that multiple cellular processes are affected, thus underpinning the complex phenotypes that arise in these cells. Based on our *in silico* analysis, nearly three quarters of the genes misregulated in *atf3*^*wt*^ mosaic EADs did not contain a consensus Atf3 site. It is therefore likely that the broad shift in the transcriptome profile also comprises an adaptive response to excess Atf3 activity. Changes to the expression of microtubule genes which translate into the alterations of the microtubule cytoskeleton represent one such example.

Importantly, the transcriptional signatures of imaginal cells overexpressing Atf3 (this study) and those lacking *scrib/dlg1* [10] significantly overlap, which could indicate common molecular and cellular mechanisms. Consistent with this notion, we identify Atf3-dependent upregulation of LamC to be a shared hallmark between the two conditions. While originally ascribed structural roles, experimental evidence link lamins to transcriptional regulation through the formation of transcriptionally repressive lamin associated domains (LADs) [66]. Thus, it is tempting to speculate that the direct and strong induction of *LamC* by Atf3 might have contributed to the wide shift in gene expression in EADs overexpressing Atf3 and/or disturbances in the microtubule network and cell polarity. However, clonal expression of a LamC transgene was insufficient to mimic defects of Atf3 overexpression or *dlg1* loss (S11 Fig), underscoring the unlikely capacity of any single target gene to recapitulate phenotypes of Atf3 gain.

Finally, it is important to note that the extent of Atf3 activity and the signaling milieu differ between *dlg1* mutant epithelia and cells overexpressing Atf3, and therefore the two conditions cannot be expected to phenocopy each other. The model presented in Figure 9 summarizes the shared and distinct roles of Atf3 in these contexts as established in this study.

## Outlook

A growing body of evidence points to striking similarities in the cellular and molecular events underlying loss of polarity and wounding [10,67]. In this context, the induction of Atf3 upon loss of polarity is in line with the early discovery of *ATF3* as a gene rapidly induced in the regenerating rat liver [68] and recent studies showing Atf3 induction during epithelial wounding [15,16]. Here we demonstrate that Atf3 levels remain low as long as epithelial polarity is intact, whereas loss of polarity due to deficiency in tumor suppressors of the Scrib complex triggers Atf3 expression. Future investigations into the link between *Drosophila* Atf3 and cell polarity are likely to unravel the impact of ATF3 expression on epithelial homeostasis and on human pathologies arising from polarity breakdown.

## MATERIALS AND METHODS

### Flies

The following fly strains were used: (a) *y w^1118^*,(b) *w*^*1118*^,(c) *y atf3*^*76*^ *w/FM7i, P{ActGFP}JMR3* [6], (d) *atf3*^*gBAC*^ [22], (e) *en-GAL4* (RRID: BDSC_30564), (f) *dpp-GAL4* (RRID: BDSC_7007), (g) *UAS-atf3*^*V*^ (this study as *atf3*^*wt*^) [6], (h) *hsFLP; act>y^+^>GAL4, UAS-GFP*, (i) *hsFLP; act>y^+^>GAL4, UAS-GFP, UAS-atf3*^*wt*^ (this study), (j) *FRT19A* (RRID: BDSC_1744), (k) *FRT82B* (RRID: BDSC_5619), (l) *UAS-p35/CyO* (RRID: BDSC_5072), (m) *GMR-hid y w FRT19; ey-Gal4, UAS-FLP* (RRID: BDSC_5248), (n) *FRT19A dlg1*^*G0342*^ (111872, DGRC), (o) *dlg1*^*m52*^ (a generous gift from F. Papagiannouli), (p) *FRT82B scrib*^*1*^ [26], (q) *eyFLP FRT19A tubGAL80; act>y^+^>GAL4*, UAS-GFP (this study), (r) *eyFLP FRT19A, tubGAL80; act>y^+^>GAL4, UAS-myrRFP* (this study), (s) *eyFLP; act>y^+^>GAL4, UAS-GFP; FRT82B, tubGAL80* [69], (t) *UAS-dlg1*^*RNAi*^ (41134, VDRC), (u) *UAS-scrib*^*RNAi*^ (105412, VDRC), (v) *TRE-DsRed* (attP16) [27], (w) *UAS-hep*^*wt*^, (x) *UAS-yki*^S111A.S168A.S250A.V5^ (RRID: BDSC_28817, this study as *yki*^*act*^), (y) *UAS-mCD8-ChRFP* (RRID:BDSC_27391, this study as *UAS-mCherry*), (z) *UAS-aPKC*^*CAAX*^ [70], (aa) *UAS-LamC* [71], (bb) *UAS-bsk*^DN^. A transgenic ATRE-GFP line was obtained using the attP16 landing site [72]. The *atf3*^*gBAC*^ fly line was used for immunostaining of Atf3::GFP. For detailed descriptions of fly crosses, see S2 Table. All crosses were carried out at 25 °C unless stated otherwise.

### Quantitative reverse transcription-PCR (qRT-PCR)

For each biological replicate, total RNA was isolated from 10 larvae or 15 adult heads with Isol-RNA Lysis Reagent (5 Prime). After DNase I treatment (Ambion, Foster City, CA), cDNA was synthesized from 1μg of RNA using oligo(dT) primers and Superscript III (Life Technologies). PCR was performed in triplicate with BioRad 2x SYBR Green mix in the CFX96 real-time PCR system (Bio-Rad, Hercules, CA). All primers were designed to anneal at 62 °C (see S1 Table for oligonucleotide sequences). All data were normalized to *rp49* transcript levels, and fold changes in gene expression were calculated using the ΔΔCt method [73].

### Genetic mosaic analysis

Clones expressing *atf3* in the wing imaginal disc were generated using a *hsFLPout* method described in [6]. Crosses were kept at 22 °C. Four days after egg lay, progeny were heat shocked for 30 min in a 37 °C water bath. Imaginal discs were dissected from wandering third instar larvae. Generation of mosaics in EADs using Mosaic Analysis with a Repressible Cell Marker method (MARCM) [74] was carried out as described in [32].

### ChIP-seq

For each genotype (see S2 Table), three biological ChIP replicates and one input replicate were generated. For each ChIP replicate, 1.2 g of adult males were collected and processed using a modified version of the protocol described in [75]. Anesthetized males were split into two batches of 600 mg and flash frozen using liquid nitrogen, pulverized with a mortar and pestle, and disrupted further via 20 strokes of a loose fitting pestle in a Dounce homogenizer containing 10 ml of Crosslinking Solution (1 mM EDTA, 0.5 mM EGTA, 100 mM NaCl, 50 mM HEPES, pH 8.0) supplemented with proteases inhibitors (11873580001, Roche). The remainder of the protocol describes the processing of a single batch. The suspension was centrifuged for 1 min at 400 g in a swing bucket rotor and then passed through two layers of 64 μm Nitex membrane into a 15 ml Falcon tube. To crosslink Atf3 to chromatin, formaldehyde was added to a final concentration of 1.8% to the Falcon tube containing 10 ml of cleared fly homogenate. The Falcon tube was then incubated on an orbital shaker at room temperature for 10 min. Crosslinking was stopped by adding glycine to a final concentration of 225 mM and shaking for a further 5 min. After centrifugation for 10 min at 1,100 g in a swing bucket rotor, cells were resuspended in 10 ml of Cell Lysis Buffer (85 mM KCl, 0.5% IGEPAL CA-630 (v/v), 5 mM HEPES, pH 8.0) supplemented with protease inhibitors and lysed in a Dounce homogenizer via twenty strokes with a tight fitting pestle. Nuclei were pelleted at 2000 g for 4 min and washed in 5 ml of Cell Lysis Buffer three times. Nuclei were resuspended in 2 ml of TBS Lysis Buffer (50 mM Tris-Cl pH 7.8, 150 mM NaCl, 1 mM EDTA pH 8.0, 1% Triton-X100, 0.01% IGEPAL CA-630) supplemented with protease inhibitors and incubated for 10 min on an orbital shaker. While submerged in an ice water bath, chromatin was sheared to 300-500 bp fragments by applying fifty 30 seconds on/30 seconds off cycles of a Branson 250 microtip sonifier with Output set to 3.5 and Duty Cycle set to 50%. The 2 ml of chromatin was split equally into two 1.5 ml low binding microcentrifuge tubes (710176, Biozym). Debris was cleared from the chromatin by centrifugation at 20,000 g for 10 min. Chromatin was cleared overnight in two tubes at 4 °C by the addition of 40 μl of sepharose-IgG beads (17-0969-01,GE) equilibrated in TBS Lysis Buffer. 100 μl of cleared chromatin was retained as an INPUT sample for sequencing. Remaining cleared chromatin was transferred to two new low binding tubes and precipitated overnight at 4 °C with 40 μl GFP Trap beads (gta-20, Chromotek) per tube. Beads were washed 5 times with cold TBS Lysis Buffer supplemented with protease inhibitors. From this point on, the beads and INPUT sample retained earlier were processed in the same fashion. Samples were resuspended in 100 μl TE buffer (1 mM EDTA, 10 mM Tris-HCl, pH 8.0) supplemented with 50 μg/ml RNase A, transferred to PCR tubes, and incubated in a thermal cycler with heated lid at 37 °C for 30 min. SDS was added to a final concentration of 0.5% (w/v) from a 10% stock proteinase K to a final concentration of 0.5 mg/ml. Tubes were then incubated at 37 °C for 12 hours followed by 65 °C for 6 h in a thermal cycler with heated lid to partially reverse the crosslinks. H_2_O was added to a final volume of 200 μl, and DNA was purified by phenol:chloroform extraction. DNA was ethanol precipitated in a low binding 1.5 ml microcentrifuge tube with sodium acetate and glycogen. DNA was washed once with 70% ethanol. DNA pellets derived from the original 1.2 g of starting adult fly material (4 microcentrifuge tubes in the end) were resuspended in 15 μl of TE.

Sequencing libraries were generated from input and ChIP samples according to the Illumina protocol for total ChIP-seq library preparation. All samples were loaded onto a single Illumina flow cell (Illumina, San Diego, CA). Using the Illumina HiSeq 2000 instrument, 50 bp of sequence were read from one DNA end. Image analysis and base calling were done with the Illumina RTA software (RRID: SCR_014332) at run time. Sequences were mapped to the *Drosophila* genome assembly (version BDGP R5/dm3, April 2006) and delivered as BAM files.

### DNA motif analyses

To identify regions bound by Atf3, BEDTools [76] (RRID: SCR_006646) was used to convert BAM files from Atf3 and control samples (3 biological ChIP replicates and one input sample each) into BED files, which were subsequently submitted to a Comparative Profile Analyzer (CoPrA) workflow to call and test the statistical significance of differentially enriched genomic locations in Atf3 ChIP-seq samples relative to control. The differential analysis depends on base-resolution coverage profiles of the Atf3 and control ChIP samples as well as the input data from both samples. They were generated from uniquely mapped reads that were extended to the original fragment length of 200 bp resulting in 6 ChIP and 2 input coverage profiles. All profiles are normalized through division of the base-wise coverage values by the total number of respective sample reads multiplied by one Million. Thereby normalized input profiles from Atf3 and control samples are subtracted from normalized Atf3 and control ChIP-profile replicates, respectively. Every processed ChIP profile replicate is then scaled separately to transform its values into the range [0,1]. They were then discretized by calculating the coverage mean of a defined sliding window. Since transcription factor binding peaks cover a small region on the genome, its size was adjusted to 50 bp. The shift is set by default to half of the window size. In the next step the discretized replicate profiles of the same sample type are combined to one resulting discretized profile by taking the mean of the window values at the same genomic position respectively. The thus processed sample profiles of Atf3 and control are finally utilized to generate a difference profile by taking the Euclidean difference between Atf3 and control values at the same genomic position. The obtained values of the difference profile are normally distributed and are located between -1 and 1 with a mean very close to 0. To filter out non-informative difference regions the profile is cleaned by removing all values within the range of 4 times the standard deviation of the mean. Remaining difference regions which are adjacent to each other and that possess the same sign are merged. They either show enrichment for Atf3 or control. They are size filtered by taking the discretization window size as a minimal threshold. The resulting difference regions are finally tested for significance by a two-sided Kolmogorov-Smirnov test of the discretized region values of Atf3 and control and multiple testing corrected by Benjamini-Hochberg. We accepted differential regions with a q-value < 0.05 that are enriched in the Atf3 sample and extracted their DNA sequences with BEDTools. Peaks were assigned to genes when located within a gene or 1 kb upstream of the transcriptional start site. Multiple genes were manually associated with a peak in the case of nested genes or dense gene regions. Enriched regions mapping to positions within the CH321-51N24 BAC (source of the *atf3*^*gBAC*^) were discarded (n=59), except for two peaks corresponding to the genes *atf3* and *CG11403*, which contained the *Drosophila* Atf3 motif.

Sequences of regions bound by Atf3 were uploaded to MEME (http://meme.nbcr.net/meme/cgi-bin/meme.cgi) for motif identification [44]. The positional weight matrix returned by MEME was submitted to TOMTOM (http://meme.nbcr.net/meme/cgi-bin/tomtom.cgi) for interspecies motif comparison [45]. FlyBase gene ontology (GO) terms identified with FlyMine (http://www.flymine.org, RRID: SCR_002694) were used for functional annotation [46].

### ChIP from eye/antennal imaginal discs

A custom protocol combining the methods described in [75,77] was used to perform ChIP against Atf3 in the EADs. For each replicate, 100 EADs with mouth hooks attached were dissected from third instar larvae in ice-cold PBS. Discs were fixed at room temperature by gently mixing in 1 ml cross-linking solution (1.8% formaldehyde, 50 mM Hepes pH 8.0, 1 mM EDTA, 0.5 mM EGTA, 100 mM NaCl), which was changed 3-4 times during fixation. Fixation was stopped by washing for 3 min in 1 ml PBS/0.01% Triton X-100/125 mM glycine 3-4 times. Fixed discs were washed for 10 min in 1 ml wash A (10 mM Hepes pH 7.6, 10 mM EDTA, 0.5 mM EGTA, 0.25% Triton X-100) and subsequently for 10 min in 1 ml wash B (10 mM Hepes pH 7.6, 200 mM NaCl, 1 mM EDTA, 0.5 mM EGTA, 0.01% Triton X-100), with the wash solutions being changed 3-4 times over the 10 min. Eye discs were separated from the brain and mouth hooks and transferred into 550 μl of TBS Lysis Buffer (see ChIP-seq protocol). Chromatin was sheared using an Active Motif EpiShear sonicator (Active Motif, 53052) equipped with a 1.5 ml EpiShear Cooled Sonication Platform (Active Motif, 53080) and 1.5 ml Benchtop Cooler (Active Motif, 53076). With Amplitude set to 50%, sonication was performed in cycles of 20 seconds on/30 seconds off for 20 minutes. Chromatin was centrifuged at 14,000 RPM for 10 min to clear debris, transferred into clean 1.5 ml tubes, and cleared overnight at 4 °C with 20 μl sepharose-IgG beads equilibrated in TBS Lysis Buffer. Cleared chromatin was transferred to new 1.5 ml tubes and precipitated overnight at 4 °C with 25 μl GFP Trap beads per tube. Beads were washed the indicated number of times with 1 ml of each of the following buffers at 4 °C on a rotating wheel for 10 min: 1 × TBS Lysis Buffer, 4 × TBS 500 Lysis Buffer (50 mM Tris-Cl pH 7.8, 500 mM NaCl, 1 mM EDTA pH 8.0, 1% Triton-X100, 0.01% IGEPAL CA-630), 1 × LiCl buffer (250 mM LiCl, 1 mM EDTA, 0.5% IGEPAL CA-630 (v/v), 10 mM Tris-HCl, pH 8.0). Beads were transferred into clean 1.5 ml tubes and washed 2 × with TE for 10 min at 4 °C on a rotating wheel. Beads were subsequently transferred into PCR tubes. Chromatin was decrosslinked and precipitated as described in the ChlP-seq protocol. Purified DNA was raised in 500 μl water.

qPCR was performed in triplicate with BioRad 2x SYBR Green mix in the CFX96 real-time PCR system (Bio-Rad, Hercules, CA). All primers were designed to anneal at 62 °C (see S1 Table 1 for oligonucleotide sequences). Data were normalized to amplification values for *ecd*. Fold enrichment of chromatin was calculated using the ΔΔC_T_ method [73].

### RNA-seq

RNA was isolated from *FRT82B atf3*^*wt*^ mosaic EADs of third instar larvae as described [78]. Total RNA libraries were generated according to the Illumina protocol and single-end sequenced on an Illumina NextSeq 500 instrument at 75 bp read length. Image analysis and base calling were done with the Illumina RTA software at run time. Published sequence data from *FRT82B* mosaic EAD samples [11] were used as control. Data were processed using a high-throughput Next-Generation Sequencing analysis pipeline [79]. Basic read quality check was performed with FastQC (v0.10.1) (RRID: SCR_014583) and read statistics were acquired with SAMtools v0.1.19 (RRID: SCR_002105) [80]. Reads were mapped to the *Drosophila* reference assembly (version BDGP R5/dm3, April 2006) using Tophat v2.0.10 (RRID: SCR_013035) [81], and gene quantification was carried out using a combination of Cufflinks v2.1.1 (RRID: SCR_014597) [82], and the DESeq2 package v1.10.1 (RRID: SCR_000154) [83], with genomic annotation from the Ensembl database (RRID: SCR_002344), version 84. In all samples, the number of total reads exceeded 50 million, from which an average 83.6 percent could be mapped, and on average 97.5% of these mapped reads fulfilled the MAPQ≥30 criterion. The results were uploaded into an in-house MySQL database and joined with BiomaRt (RRID: SCR_002987) v2.26.1 [84] annotations from Ensembl, version 84. Lists of differentially expressed genes were defined by a final database export using 5 and 0.01 as cutoffs for DESeq2-based FCs and p-values, respectively. To identify genes differentially expressed under the respective conditions, the average of at least three biological replicates was calculated. The S3 Dataset shows all transcripts whose expression differed ≥ 1.5-fold in *FRT82B atf3*^*wt*^ compared to *FRT82B* control.

### Atf3 motif search

The experimentally derived *Drosophila* Atf3 PWM was submitted to the online FIMO utility (http://meme-suite.org/tools/fimo) (RRID: SCR_001783) to identify Atf3 motifs in *Drosophila melanogaster* genome (UCSC, dm3), with a p-value threshold set at 0.0001. FIMO results, in the form of a BED file, were subsequently submitted to the online PAVIS tool (http://manticore.niehs.nih.gov/pavis3), using default settings.

### Plasmid constructs

To create the Atf3 expression reporter, oligonucleotides containing four intact or mutated Atf3 sites (see S1 Table for sequences) were cloned via Mlu1 and Not1 sites into the pRedRabbit and pGreenRabbit vectors [85]. To express N-terminally tagged GFP- or FLAG-Atf3 proteins from the UAST promoter, *atf3* cDNA (see S1 Table for oligonucleotide sequences) was cloned into pENTR4 and subsequently recombined using LR Clonase II (11791-020, Life Technologies) into pTGW and pTFW vectors, respectively (T. Murphy, *Drosophila* Genomic Resource Center).

### S2 cell culture

Schneider 2 (S2) cells were cultured at 25 °C in Shields and Sang M3 insect medium (S8398-1L, Sigma-Aldrich) containing 8% fetal bovine serum (Gibco, Life Technologies) without antibiotics. Cells were transfected using X-tremeGENE (Roche Applied Science). Expression of UAS-driven genes was induced by co-transfection with a pWA-GAL4 plasmid expressing GAL4 under an *actin5C* promoter. Cells were fed 24 hours after transfection. Cells were imaged or lysed 72 hours after transfection.

### Tissue staining

Tissues from third instar larvae were processed as described previously [6]. The following primary and secondary antibodies were used at the indicated dilutions: anti-Dlg1 (RRID: AB_528203; 1:200), anti-Fasciclin III (RRID: AB_528238; 1:200), anti-Elav (RRID: AB_528217 and AB_528217; 1:200), anti-DE-cad (RRID: AB_528120; 1:200), anti-Crumbs (RRID: AB_528181; 1:200), anti-p120-catenin (RRID: AB_2088073; 1:200), anti-α–Tubulin (RRID: AB_579793; 1:200), anti-β–Tubulin (RRID: AB_2315513; 1:200), anti-LaminC (RRID: AB_528339; 1:500) and anti-βPS integrin (RRID: AB_528310; 1:200) from the Developmental Studies Hybridoma Bank (DSHB) (Iowa City, Iowa), rabbit-anti γ-Tubulin (T6557, Sigma Aldrich; 1:50), rabbit anti-GFP (G10362, Invitrogen; 1:500), rabbit-anti-Rab5 (ab31261, Abcam; 1:200), mouse-anti-Rab11 (BD610656, BD Biosciences; 1:200), chicken-anti-Avl (a gift from D. Bilder, 1:500) and Cy2, Cy3- and Cy5-conjugated secondary antibodies (Jackson Immunoresearch). Tissues were counterstained with DAPI (1 μg/ml) and mounted in DABCO-Mowiol medium (Sigma-Aldrich).

### Dextran uptake assay

Third instar imaginal discs were incubated in M3 media containing 0.5 mM MW3000 Texas Red dextran (D3328; ThermoFisher Scientifc) for 15 minutes (pulse) at 25 °C. Discs were briefly washed 2 times in M3 media followed by incubation in M3 media for 60 minutes (chase). Discs were subsequently fixed in PBS-1% formaldehyde for 20 min, counterstained with DAPI, and mounted in DABCO-Mowiol medium (Sigma-Aldrich).

### Image acquisition and processing

S2 cells were imaged using a DP72 camera mounted on an Olympus CK4X41 miscroscope, in conjunction with cellSens 1.1 Software (RRID: SCR_014551). Confocal images were acquired at room temperature using an Olympus FV1000 confocal microscope equipped with 20x UPlan S-Apo (NA 0.85), 40x UPlan FL (NA 1.30) and 60x UPlanApo (NA 1.35) objectives. Transversal sections were generated using Imaris 7.0.0 (Bitplane) (RRID: SCR_007370). Figure assembly and image brightness and contrast adjustments were done in Photoshop CS5.1 (Adobe Systems, Inc.) (RRID: SCR_014199). Z-stacks of adult eyes were taken using a motorized Leica M165 FC fluorescent stereomicroscope equipped with the DFC490 CCD camera. Images were processed using the Multifocus module of LAS 3.7.0 software (Leica).

### Flow cytometry of eye/antenna imaginal discs

Third instar larvae were collected and washed 2x with PBS. EADs were dissected in PBS (on average sixty EADs /replicate/genotype) and transferred to 1.5 ml low binding microcentrifuge tubes (no more than one hour before dissociation). PBS was removed from EADs and was replaced with 100 μl dissociation solution containing 1 mg/ml collagenase I (Sigma, C2674), and 1 mg/ml papain (Sigma P4762). Samples were incubated for 60 min at room temperature and gently swirled every 15 min. After the dissociation solution was removed, discs were carefully rinsed with 500 μl PBS, which was then replaced with 100 μl of PBS. The final dissociation was performed by passing the discs through a 27G insulin syringe (Terumo) five times. Additional 200 μl of PBS was added for a final volume of 300 μl (5 μl PBS per disc). Following sample filtration through a Filcone filter, propidium iodide was added to measure cell viability and samples were stored on ice until sorting. For each sample (n≥3 per genotype), 30,000 events were counted on a BD LSRFortessa Cell Analyzer in combination with BD FACSDiva software v8.0 (both BD Bioscience) using gates set to distinguish GFP+, GFP-, and PI-cells.

### Scanning electron microscopy

Adult heads were fixed in 80% ethanol, postfixed with 1% osmium tetroxide, dehydrated in ethanol, critical point dried, gold coated, and observed under a JEOL JSM-7401F (Tokyo, Japan) 6300 scanning electron microscope.

### Immunoblotting

S2 cells were lysed in 50 mM Tris-HCl (pH 7.8), 150 mM NaCl, 1 mM EDTA (pH 8.0), 1% Triton X-100, 0.01% Igepal, and protease inhibitors (Roche Applied Science). Protein concentration was quantified using Pierce 660 reagent (22660, Thermo Scientific) according to manufacturer’s instructions. Following SDS-PAGE, proteins were detected by immunoblotting with mouse anti-Flag M2 (1:1000, Sigma Aldrich), rabbit anti-GFP (1:5000, TP401, Acris) and mouse anti-α-Spectrin (1:1000, RRID: AB_528473) antibodies, followed by incubation with corresponding HRP-conjugated secondary antibodies (Jackson Immuno Research). Chemiluminescent signal was captured using ImageQuant LAS4000 reader (GE Healthcare).

### Sample Size Criteria

For sample size criteria, *post hoc* analysis of results presented in Figs 1, S1, S3, S5 and S9 using G*Power 3.1 (RRID: SCR_013726) [86] determined that the statistical power of all statistically significant differences (1-β) exceeded 0.94, given sample size, mean, and standard deviation for each condition.

## Acknowledgements

We thank Dirk Bohmann, Sarah Bray, Sonsoles Campuzano, Fani Papagiannouli, Andreas Wodarz, Lori Wallrath, David Bilder, the Bloomington Stock Center (Bloomington, USA), the Vienna *Drosophila* Resource Center (VDRC, Vienna, Austria), NIG-Fly Stock Center (Mishima, Japan), the Developmental Studies Hybridoma Bank (DSHB, Iowa, USA) and the *Drosophila* Genomics Resource Center (Bloomington, USA) for plasmids, fly stocks, and antibodies. We are grateful to Alvaro Rada-Iglesias for advice on ChIP, Vladimir Benes and the Genomics Core Facility in EMBL for ChIP and mRNA sequencing and Peter Frommolt and Prerana Wagle from the CECAD Bioinformatics Facility Cologne for mRNA-seq data processing, and Merve Kilinc for assistance with FACS. We acknowledge the Laboratory of Electron Microscopy (Biology Center, CAS), supported by the Czech-BioImaging large RI project LM2015062. We thank Tina Bresser and Nils Teuscher for technical assistance.

## Author contribution

C.D., A.C., G.C. and M.U. conceived the study, designed and performed the experiments, and analyzed the data. M.J. prepared the SEM samples. B.H and C.Kl. developed and described the CoPrA workflow. C.D., C.Kl., and B.H. analyzed the ChIP-seq data. C.D., G.C., M.U., and M.J. wrote the paper. M.U. acquired funding.

## Supplementary material

**S1 Fig. The *dlg1*^*G0342*^ mutation is a loss-of-function allele**

**(A)** No *dlg1* transcript is detected in hemizygous *dlg1*^*G0342*^ larvae relative to control. **(B-E)** *eyFLP*-mediated mitotic recombination was used to generate clones (GFP) in the EADs of the indicated genotypes. Homozygous *dlg1*^*G0342*^ and *atf3*^*76*^ *dlg1*^*G0342*^ EAD clones (B-C and D-E, respectively) are deficient for Dlg1 protein (B’-E’). qRT-PCR data are means of 6 biological replicates. Error bars depict 95% confidence interval; Unpaired Student’s t-tests assuming unequal variance were used to calculate p-values: ***= p<0.001. Scale bars: 100 μm (B,D), 10 μm (C,E).

**S2 Fig. Atf3 activates an ATRE reporter in cell culture**

**(A,B)** In S2 cells, Atf3 induced the ATRE-RFP reporter compared to undetectable activity in control cells. **(C,D)** Atf3 did not activate a mutant mATRE-RFP reporter. **(E-G)** Expression of FLAG::Atf3 strongly induced an ATRE-GFP reporter in S2 cells. While co-expression of Hep/JNKK and Bsk/JNK induced an AP-1 reporter (TRE-DsRed) (F,G), it only weakly activated the ATRE reporter (E). *α*-Spectrin served as a loading control.

**S3 Fig. Atf3 drives specific phenotypes in *dlg1* null mutant mosaic eyes**

**(A-G)** *eyFLP*-mediated mitotic recombination was used to generate clones of the indicated genotypes in the eye. Homozygous *dlg1*^*G0342*^ or *dlg1*^*m52*^ EAD clones lead to smaller adult eyes of irregular shape (F,G) containing patches of undifferentiated tissue (A,D) relative to control (Fig 2A). Adult eyes derived from *atf3*^*76*^ *dlg1*^*G0342*^ or *atf3*^*76*^ *dlg1*^*m52*^ mosaic EADs exhibit mild or no differentiation defects (B,E) as well as restored eye size and shape (F,G). A single copy of a genomic *atf3*^*gBAC*^ reinstates differentiation defects to *atf3*^*76*^ *dlg1*^*G0342*^ adult eyes (C). Outlines of adult eyes from the indicated genotypes are presented vertically aligned along their midline (G). Adult eye measurements were performed from at least 17 biological replicates. Error bars depict 95% confidence interval; Unpaired Student’s t-tests assuming unequal variance were used to calculate p-values: ***= p<0.001 ****p<0.0001.

**S4 Fig. Clonal expression of *atf3* does not impact differentiation in the EAD**

**(A-D)** *eyFLP*-mediated mitotic recombination was used to generate clones (GFP) in the EAD of the indicated genotypes. *dlg1*^*G0342*^ mutant cells located to the left of the morphogenetic furrow (arrow) are often Elav-negative (A,B) compared to a regular Elav staining pattern in the *atf3*^*wt*^ clones (C,D) and the non-clonal neighbors. Discs were counterstained with DAPI. Micrographs are single confocal slices. All images show EADs 7 days after egg laying. Scale bars: 100 μm (A,C), 20 μm (B,D).

**S5 Fig. JNK signaling is elevated upon loss of polarity and blocking apoptosis does not mitigate the genetic interaction between Atf3 and Dlg1**

**(A-H)** *eyFLP*-mediated mitotic recombination was used to generate GFP-labeled clones of the indicated genotypes in the EAD. **(A)** Flow cytometry determined the number of GFP+ cells in EADs bearing clones of the indicated genotypes. Relative to control (n=4), *dlg1*^*G0342*^ (n=3), *atf3*^*76*^ *dlg1*^*G0342*^ (n=4), and *dlg1*^*G0342*^ *atf3*^*wt*^ (n=5) clones were less abundant. *dlg1*^*G0342*^ *atf3*^*wt*^ cells were less frequent relative to *dlg1*^*G0342*^ alone. Blocking apoptosis raised the relative abundance of *dlg1*^*G0342*^ (n=7) and *atf3*^*76*^ *dlg1*^*G0342*^ (n=6) cells, but not to control (n=4) levels. Unpaired Student’s t-tests assuming unequal variance were used to calculate p-values. Error bars reflect the 95% confidence interval. ***= p<0.001. **(B-C)** An AP-1 reporter (TRE-DsRed) serves as a readout of JNK pathway activity and is upregulated in *dlg1*^*G0342*^ (B’) and *atf3*^*76*^*dlg1*^*G0342*^ (C’) EAD clones. **(D)** The eclosion rate of animals bearing *atf3*^*76*^*dlg1*^*G0342*^*p35* EAD was less than control but was four times higher than animals bearing *dlg1*^*G0342*^*p35* EAD. Four biological replicates were used for each genotype. Unpaired Student’s t-tests assuming unequal variance were used to calculate p-values: ***= p<0.001. **(E-H)** Compared to control (E), adult eyes derived from *dlg1*^*G0342*^*p35* EAD were very small, comprising mostly undifferentiated tissue (F), while those derived from *atf3*^*76*^*dlg1*^*G0342*^*p35* EAD exhibited only traces (G) or small patches (H) of defective photoreceptor differentiation. **(I-K)** The EGUF/hid technique was used to generate adult eyes comprised entirely of control (I), *dlg1*^*G0342*^ (J) and *atf3*^*76*^*dlg1*^*G0342*^ (K) clonal tissue as non-clonal cells were removed by expression of pro-apoptotic protein Hid. Compared to control (I), *dlg1*^*G0342*^ eyes were severely reduced in size with only traces of ommatidia left (J). *atf3*^*76*^*dlg1*^*G0342*^ eyes were only mildly reduced relative to control and contained several rows of orderly arranged ommatidia (K). Micrographs (B,C) are projections of multiple confocal sections showing EADs 7 days after egg laying. Scale bars: 100 μm (B,C).

**S6 Fig. Epithelial clones lacking *dlg1* show Atf3-dependent perturbation of early endosomes, but no defects in endocytosis**

**(A-F)** *eyFLP*-mediated mitotic recombination was used to generate clones (GFP) in the EADs of the indicated genotypes. Apoptosis of mutant cells was reduced by p35 overexpression. **(A-C)** *dlg*^*G03421*^ and *atf3*^*76*^*dlg1*^*G0342*^ clones show no disturbances in the uptake of fluorescently labeled dextran (B’- C’) compared to control clones (A’). **(D-E)** The regular pattern of the early endosomal marker Avl (D’) is disrupted in *dlg1* mutant cells (E’), but restored in *atf3*^*76*^*dlg1*^*G0342*^ double mutant clones (F’). Discs were counterstained with DAPI. White arrows indicate cross sections, which appear below the corresponding panels and are oriented apical side up. Clones are outlined by white dotted lines. All images show EAD 7 days after egg laying. Scale bars: 10 μm (A-F).

**S7 Fig. Removal of *atf3* restores distribution of Rab5-positive vesicles in *dlg1* mutant clones**

**(A-F)** *eyFLP*-mediated mitotic recombination was used to generate clones (GFP) in the EADs of the indicated genotypes. Lower levels and disturbed pattern of the early endosome marker Rab5 (B’,E’) observed in *dlg1*^*G0342*^ clones relative to surrounding and control tissue (A’,D’) were largely restored in *atf3*^*76*^*dlg1*^*G0342*^ clones (C’,F’). Discs were counterstained with DAPI. In A-C, EADs and in D’-F’, clones are outlined with dotted white lines. White arrows indicate cross sections, which appear below the corresponding panels and are oriented apical side up. On the cross sections of D’-F’, arrowheads indicate the regular Rab5 accumulation in the photoreceptors. Micrographs are projections of multiple confocal slices (A-F). Scale bars: 100 μm (A-C), 20 μm (D-F).

**S8 Fig. Gain of Atf3 alters cellular trafficking machinery and the localization of polarity determinants**

**(A-B)** Abundance of Rab5 vesicles (A’) was reduced in cells expressing Atf3 (*hsFLPout>>atf3*^*wt*^) (GFP), compared to surrounding tissue while the amount of Rab11 endosomes increased and showed a distinct basal accumulation in Atf3 overexpressing cells (*en>atf3*^*wt*^) (GFP) compared to wild type epithelium (B’). **(C-D)** Cross sections of immunostained wing imaginal disc clones expressing Atf3 (*hsFLPout>>atf3*^*wt*^) revealed lower levels of the polarity determinant Crumbs on the apical surface (C’). In contrast, the βPS integrin Myospheroid (Mys) appeared to be more abundant in Atf3 overexpressing cells and was distributed along the lateral membranes (D’). White dotted lines indicate clones (A,C,D) or the posterior compartment (B) in the wing disc. Arrowheads indicate cross sections, which appear below the corresponding panels and are oriented apical side up. Micrographs are otherwise single confocal slices. Scale bars: 10 μm (A-D).

**S9 Fig. Atf3 binds to ATRE sites in the LamC gene and induces LamC transcription in larval EADs**

**(A)** Independent qRT-PCR from newly isolated mosaic EADs confirmed RNA-seq results showing enrichment of *LamC, dynactin, ude* and *arp1* transcripts in EADs overexpressing Atf3 (n=3) relative to control (n=3) while *alphaTub84B* and *betaTub56D* remained unchanged. **(B)** ChIP from third instar EADs followed by qPCR was used to quantify enrichment of the indicated Atf3 sites in *atf3*^*gBAC*^ samples (n=3) relative to *w*^*1118*^ control (n=3). *LamC* and *α-Tub84B* were enriched in *atf3*^*gBAC*^ samples compared to control. Note that *LamC* was bound by Atf3 at two of the three Atf3 sites within the gene. Atf3 did not bind the ATRE site in *coracle* which was occupied in samples from adults. Error bars indicate 95% confidence interval; Unpaired Student’s t-tests assuming unequal variance were used to calculate p-values: **p=0.007 (*LamC*), *p=0.018 (*dynactin*), **p=0.008 (*ude*), *p=0.024 (*arp1*).

**S10 Fig. Transcriptional response to gain of Atf3 overlaps with genes misregulated by depletion of the Scrib polarity module**

**(A-B)** Venn diagrams show the overlap between genes differentially regulated by a factor ≥1.5 in mosaic EADs overexpressing Atf3 (*eyFLP*^*MARCM*^ *atf3*^*wt*^) and wing imaginal discs of *scrib*^*1*^ homozygous or *dlg*^*40-2*^/*Y* hemizygous mutant larvae. Contingency tables provide information about directionality of expression of shared transcripts between the two datasets and serve to calculate the significance of overlap using a one-tailed Fisher Exact Probability test.

**S11 Fig. LamC overexpression in imaginal disc cells is not sufficient to recapitulate Atf3 gain-of-function phenotypes**

**(A-B)** Immunostaining for α-Tubulin (A) and Crumbs (B) is unchanged in wing imaginal disc clones (GFP) expressing wild type LamC (*hsFLPout>>LamC*). Discs were counterstained with DAPI. Clones are outlined with dotted white lines. White arrows indicate cross sections, which appear below the corresponding panels and are oriented apical side up. Scale bars: 10 μm (A,B).

**S1 Table. Summary of oligonucleotides**

**S2 Table. Summary of fly crosses**

**S1 Dataset. Atf3 ChIP-seq peaks and targets**

**(A)** Dataset All Peaks (n=152). The chromosomal positions (Chromosome, Peak Start, Peak End) of the 152 regions differentially enriched in Atf3 ChIP samples are given.

**(B)** Dataset Atf3 Peaks (n=112). Results from MEME (Bailey and Elkan, 1994), identifying a consensus Drosophila Atf3 motif in 112 peaks. For each peak containing the *Drosophila* Atf3 motif, the Start Position and End Position are given, and the probability of a random string having the same MEME match score or higher (p-value) is also provided. The core twelve nucleotide sequence (Atf3 Motif Sequence) and the immediate upstream and downstream sequence (5' Flanking Sequence, 3' Flanking Sequence) of each Atf3 motif are given. Genes most proximal to the Atf3 motifs are provided as FlyBase identifiers (Associated Genes (Symbols) and Associated Genes (FlyBase ID)).

**(C)** Dataset Cytoskeleton GO. Genes belonging to at least one of five cytoskeleton gene ontology (GO) terms (actin binding GO_0003779, calponin homology domain IPR_001715, cytoskeletal protein binding GO_0008092, cytoskeleton organization GO_0007010, cytoskeleton GO_0005856) are listed with their CG Number, Gene Symbol, and FlyBase ID. GO terms are arranged in columns. Genes belonging to a given GO term are indicated with green cells containing an “x’’.

**S2 Dataset. *Drosophila* Atf3 PWM**

**S3 Dataset. Transcriptomic analysis of mosaic EADs**

